## Supplemental material for "Metabolism of ʟ -arabinose converges with virulence regulation to promote enteric pathogen fitness"

**Supplementary information:**

Table S1 – Summary of differentially expressed genes identified by RNA-seq
Table S2 – Bacterial strains used in this study
Table S3 – Plasmids used in this study
Table S4 – Primers used in this study
Supplementary refernces

**Supplementary Table 1** – Summary of differentially expressed genes identified by RNA-seq. Data derived from EHEC cultures grown in MEM-HEPES alone (control) or supplemented with L-arabinose (treatment).

| Feature ID | Fold change | Log2 FC | FDR *p*-value | Function |
|---|---|---|---|---|
| ***Upregulated*** | | | | |
| *araH* | 151.81 | 7.25 | 3.49E-136 | High-affinity L-arabinose transport system |
| *ygeA* | 149.37 | 7.22 | 1.08E-126 | Amino acid racemase |
| *araF* | 145.09 | 7.18 | 5.17E-137 | L-arabinose ABC transporter periplasmic binding |
| *araE* | 139.81 | 7.13 | 1.39E-131 | L-arabinose:H+ symporter |
| *araG* | 126.13 | 6.98 | 1.42E-132 | L-arabinose ABC transporter ATP binding subunit |
| *araD* | 98.47 | 6.62 | 2.16E-109 | L-ribulose-5-phosphate 4-epimerase |
| *araA* | 59.08 | 5.88 | 4.47E-107 | L-arabinose isomerase |
| *lysR* | 46.53 | 5.54 | 6.61E-89 | DNA-binding transcriptional dual regulator |
| *Z0417* | 43.15 | 5.43 | 6.77E-75 | Putative ATP-binding component of transport system |
| *araB* | 42.78 | 5.42 | 3.31E-98 | L-ribulokinase |
| *Z0415* | 26.51 | 4.73 | 1.55E-79 | Putative periplasmic binding protein |
| *ninG* | 21.44 | 4.42 | 0.04 | Unknown protein encoded by prophage CP-933K |
| *araJ* | 12.75 | 3.67 | 1.19E-38 | Putative transport protein |
| *rutE* | 10.91 | 3.45 | 2.01E-32 | Putative malonic semialdehyde reductase |
| *rutF* | 9.22 | 3.20 | 4.27E-29 | FMN reductase |
| *ddpB* | 9.04 | 3.18 | 5.00E-14 | Putative D,D-dipeptide ABC transporter subunit |
| *rutA* | 8.32 | 3.06 | 4.40E-30 | Pyrimidine monooxygenase |
| *nac* | 8.1 | 3.02 | 9.50E-30 | DNA-binding transcriptional dual regulator |
| *cbl* | 8.08 | 3.01 | 2.51E-29 | DNA-binding transcriptional activator |
| *rutB* | 8.05 | 3.01 | 5.29E-28 | Ureidoacrylate amidohydrolase |
| *rutD* | 7.64 | 2.93 | 9.66E-27 | Putative aminoacrylate hydrolase |
| *rutC* | 7.53 | 2.91 | 1.21E-25 | 3-aminoacrylate deaminase |
| *glnK* | 7.4 | 2.89 | 6.52E-27 | Nitrogen regulatory protein PII-2 |
| *yegT* | 7.35 | 2.88 | 2.31E-15 | Putative nucleoside permease protein |
| *rutG* | 7.31 | 2.87 | 8.10E-26 | Pyrimidine:H+ symporter |
| *yegU* | 7.09 | 2.83 | 1.13E-15 | Putative aminoacrylate hydrolase |
| *Z0418* | 7 | 2.81 | 1.77E-19 | Putative permease component of transport system |
| *yegV* | 6.87 | 2.78 | 1.03E-16 | Putative sugar kinase |
| *epd* | 6.82 | 2.77 | 2.46E-24 | D-erythrose-4-phosphate dehydrogenase |
| *yjfF* | 6.66 | 2.74 | 6.61E-25 | Galactofuranose ABC transporter putative subunit |
| *amtB* | 6.58 | 2.72 | 1.21E-24 | Ammonium transporter |
| *Z4629* | 6.35 | 2.67 | 4.68E-21 | Putative periplasmic binding transport protein |
| *glnA* | 6.23 | 2.64 | 1.74E-23 | Glutamine synthetase |
| *sgrT* | 5.95 | 2.57 | 1.50E-17 | Glucose uptake inhibitor |
| *yrbN* | 5.86 | 2.55 | 4.18E-03 | Hypothetical protein |
| *Z3167* | 5.36 | 2.42 | 0.04 | CP4-44 prophage; putative uncharacterised protein |
| *yneE* | 5.24 | 2.39 | 3.30E-19 | PF01062 family inner membrane protein |
| *ytfT* | 5.2 | 2.38 | 2.13E-19 | Galactofuranose ABC transporter putative subunit |
| *yidA* | 5.12 | 2.36 | 3.93E-16 | Sugar phosphatase |
| *galP* | 5.01 | 2.32 | 2.81E-16 | Galactose:H+ symporter |
| *cheB* | 5.01 | 2.32 | 0.02 | Glutamate methylesterase/ glutamine deamidase |
| *ddpX* | 4.9 | 2.29 | 1.78E-15 | D-alanyl-D-alanine dipeptidase |
| *ytfR* | 4.87 | 2.28 | 4.10E-18 | Galactofuranose ABC transporter ATP binding subunit |
| *ydiH* | 4.29 | 2.10 | 1.43E-13 | Hypothetical protein |

| | | | | |
|---|---|---|---|---|
| *malE* | 4.23 | 2.08 | 4.20E-14 | Maltose ABC transporter periplasmic binding protein |
| *hdeB* | 4.23 | 2.08 | 2.25E-13 | Periplasmic acid stress chaperone |
| *ompC_1* | 4.14 | 2.05 | 7.98E-15 | Outer membrane porin C |
| *malF* | 3.9 | 1.96 | 3.45E-12 | Maltose ABC transporter membrane subunit |
| *gltJ* | 3.9 | 1.96 | 1.31E-11 | Glutamate/aspartate ABC transporter subunit |
| *malK* | 3.83 | 1.94 | 6.34E-11 | Maltose ABC transporter ATP binding subunit |
| *fbp* | 3.79 | 1.92 | 4.05E-13 | Fructose-1,6-bisphosphatase 1 |
| *Z0414* | 3.7 | 1.89 | 1.45E-08 | Hypothetical protein |
| *yedL* | 3.68 | 1.88 | 6.34E-11 | Putative acetyltransferase |
| *tam* | 3.63 | 1.86 | 2.00E-12 | *Trans*-aconitate 2-methyltransferase |
| *xylF* | 3.63 | 1.86 | 3.54E-11 | Xylose ABC transporter periplasmic binding protein |
| *Z2983* | 3.62 | 1.86 | 2.40E-10 | Putative tail fiber assembly protein of CP-933T |
| *nudK* | 3.61 | 1.85 | 1.61E-07 | GDP-mannose hydrolase |
| *fhuD* | 3.61 | 1.85 | 4.62E-07 | Iron(III) hydroxamate ABC transporter |
| *gadC* | 3.59 | 1.84 | 3.36E-12 | L-glutamate:4-aminobutyrate antiporter |
| *tusC* | 3.58 | 1.84 | 8.12E-08 | Sulphurtransferase complex subunit |
| *adhE* | 3.54 | 1.82 | 4.76E-12 | Fused acetaldehyde-CoA dehydrogenase |
| *glnL* | 3.53 | 1.82 | 1.54E-10 | Protein histidine kinase |
| *glnH* | 3.46 | 1.79 | 2.76E-11 | L-glutamine ABC transporter periplasmic binding |
| *Z2981* | 3.41 | 1.77 | 2.74E-06 | IS629 transposase encoded within prophage CP-933T |
| *Z2377* | 3.41 | 1.77 | 0.04 | Unknown protein encoded within prophage CP-933R |
| *bioF* | 3.37 | 1.75 | 4.45E-10 | 8-amino-7-oxononanoate synthase |
| *purA* | 3.27 | 1.71 | 1.65E-10 | Adenylosuccinate synthetase |
| *Z1539* | 3.25 | 1.70 | 6.57E-08 | Hypothetical protein |
| *glnP* | 3.24 | 1.70 | 2.99E-09 | L-glutamine ABC transporter membrane subunit |
| *lsrG* | 3.23 | 1.69 | 1.69E-10 | (4*S*)-4-hydroxy-5-phosphonooxypentane-2,3-dione |
| *avtA* | 3.21 | 1.68 | 9.09E-09 | Valine-pyruvate aminotransferase |
| *ruvX* | 3.21 | 1.68 | 5.77E-06 | Holliday junction resolvase |
| *rimL* | 3.17 | 1.66 | 3.00E-09 | 50S ribosomal protein L7/L12-serine acetyltransferase |
| *Z2976* | 3.16 | 1.66 | 3.02E-07 | Unknown protein encoded by prophage CP-933T |
| *lsrF* | 3.12 | 1.64 | 5.56E-10 | 3-hydroxy-2,4-pentadione 5-phosphate thiolase |
| *lamB* | 3.1 | 1.63 | 3.93E-09 | Maltose OM channel/phage lambda receptor |
| *Z5892* | 3.09 | 1.63 | 3.04E-07 | Hypothetical protein |
| *galS* | 3.06 | 1.61 | 1.63E-09 | DNA-binding transcriptional dual regulator |
| *fruB* | 3.05 | 1.61 | 4.05E-09 | Fructose-specific PTS multiphosphoryl transfer protein |
| *gltI* | 3.04 | 1.60 | 1.64E-09 | Glutamate/aspartate ABC transporter periplasmic |
| *bioC* | 2.98 | 1.58 | 1.71E-06 | Malonyl-acyl carrier protein methyltransferase |
| *ddpA* | 2.97 | 1.57 | 2.29E-08 | Putative D,D-dipeptide ABC transporter periplasmic |
| *nfo* | 2.97 | 1.57 | 3.91E-06 | Endonuclease IV |
| *yobD* | 2.95 | 1.56 | 3.45E-03 | DUF986 domain-containing inner membrane protein |
| *ptsP* | 2.94 | 1.56 | 4.65E-08 | Phosphoenolpyruvate-protein phosphotransferase |
| *glnG* | 2.92 | 1.55 | 3.04E-08 | DNA-binding transcriptional dual regulator |
| *cspG* | 2.91 | 1.54 | 6.09E-07 | Cold shock protein |
| *cytR* | 2.9 | 1.54 | 5.33E-08 | DNA-binding transcriptional repressor |
| *galT* | 2.89 | 1.53 | 1.38E-08 | Galactose-1-phosphate uridylyltransferase |
| *aroG* | 2.87 | 1.52 | 3.20E-08 | 3-deoxy-7-phosphoheptulonate synthase |
| *nlpI* | 2.86 | 1.52 | 1.44E-08 | Lipoprotein |
| *wcaF* | 2.86 | 1.52 | 0.02 | Colanic acid biosynthesis acetyltransferase |
| *ydiQ* | 2.86 | 1.52 | 0.05 | Putative electron transfer flavoprotein subunit |
| *lsrK* | 2.83 | 1.50 | 1.91E-08 | Autoinducer-2 kinase |
| *galK* | 2.83 | 1.50 | 2.60E-08 | Galactokinase |

| | | | | |
|---|---|---|---|---|
| *trmL* | 2.82 | 1.50 | 5.13E-05 | tRNA (cytidine/uridine-2'-O)-ribose methyltransferase |
| *gmm* | 2.81 | 1.49 | 0.03 | GDP-mannose mannosyl hydrolase |
| *yijF* | 2.8 | 1.49 | 0.01 | DUF1287 domain-containing protein |
| *hutW* | 2.75 | 1.46 | 4.77E-07 | Putative coproporphyrinogen III oxidase |
| *dmsD* | 2.74 | 1.45 | 4.96E-04 | Redox enzyme maturation protein |
| *gltL* | 2.73 | 1.45 | 7.78E-07 | Glutamate/aspartate ABC transporter |
| *hda* | 2.69 | 1.43 | 1.27E-06 | Inibitor of reinitiation of DNA replication |
| *lsrR* | 2.68 | 1.42 | 1.20E-07 | DNA-binding transcriptional repressor |
| *cydD* | 2.67 | 1.42 | 3.42E-06 | Glutathione/L-cysteine ABC exporter subunit |
| *phrB* | 2.66 | 1.41 | 7.59E-07 | Deoxyribodipyrimidine photo-lyase |
| *fhuC* | 2.66 | 1.41 | 1.10E-05 | Iron(III) hydroxamate ABC transporter |
| *nei* | 2.66 | 1.41 | 1.05E-04 | Endonuclease VIII |
| *Z2972* | 2.64 | 1.40 | 2.17E-07 | Unknown protein encoded by prophage CP-933T |
| *bglJ* | 2.64 | 1.40 | 5.16E-06 | DNA-binding transcriptional regulator |
| *rstA* | 2.64 | 1.40 | 2.39E-04 | DNA-binding transcriptional regulator |
| *ygaH* | 2.64 | 1.40 | 1.31E-03 | L-valine exporter |
| *Z_RS32400* | 2.64 | 1.40 | 2.79E-03 | Hypothetical protein |
| *Z4628* | 2.64 | 1.40 | 0.01 | Membrane protein |
| *bioD* | 2.62 | 1.39 | 1.06E-04 | Dethiobiotin synthetase |
| *fau* | 2.61 | 1.38 | 1.32E-05 | Putative 5-formyltetrahydrofolate cyclo-ligase |
| *ytfQ* | 2.6 | 1.38 | 2.85E-07 | Galactofuranose ABC transporter periplasmic binding |
| *ydfH* | 2.6 | 1.38 | 1.09E-05 | DNA-binding transcriptional repressor |
| *btsR* | 2.6 | 1.38 | 1.02E-04 | DNA-binding transcriptional dual regulator |
| *mglB* | 2.56 | 1.36 | 4.64E-07 | D-galactose/methyl-galactoside ABC transporter |
| *yoaG* | 2.56 | 1.36 | 4.48E-03 | DUF1869 domain-containing protein |
| *ybaQ* | 2.53 | 1.34 | 2.62E-06 | Hypothetical protein |
| *malG* | 2.53 | 1.34 | 1.01E-05 | Maltose ABC transporter membrane subunit |
| *yigA* | 2.53 | 1.34 | 1.33E-04 | DUF484 domain-containing protein |
| *ydhX* | 2.53 | 1.34 | 0.02 | Putative 4Fe-4S ferredoxin-like protein |
| *glnQ* | 2.52 | 1.33 | 1.00E-05 | L-glutamine ABC transporter ATP binding subunit |
| *galE* | 2.51 | 1.33 | 1.03E-06 | UDP-glucose 4-epimerase |
| *ptsN* | 2.51 | 1.33 | 1.77E-05 | Phosphotransferase system enzyme IIA |
| *yfiB* | 2.5 | 1.32 | 3.00E-04 | Lipoprotein |
| *ymcF* | 2.5 | 1.32 | 3.29E-04 | Hypothetical protein |
| *yggU* | 2.49 | 1.32 | 0.01 | DUF167 domain-containing protein |
| *Z1538* | 2.48 | 1.31 | 5.12E-06 | Putative pilin |
| *lipB* | 2.48 | 1.31 | 8.08E-05 | Lipoate biosynthesis protein |
| *yfaE* | 2.47 | 1.30 | 0.02 | Ferredoxin-like diferric-tyrosyl radical cofactor protein |
| *nhaR* | 2.45 | 1.29 | 1.08E-05 | DNA-binding transcriptional activator |
| *Z2970* | 2.44 | 1.29 | 2.14E-06 | Putative regulator for prophage CP-933T |
| *kduD* | 2.44 | 1.29 | 1.13E-05 | Putative 2-keto-3-deoxy-D-gluconate dehydrogenase |
| *fkpB* | 2.44 | 1.29 | 1.16E-03 | Peptidyl-prolyl cis-trans isomerase |
| *argO* | 2.44 | 1.29 | 7.57E-03 | L-arginine exporter |
| *bioA* | 2.43 | 1.28 | 4.24E-06 | Adenosylmethionine-8-amino-7-oxo aminotransferase |
| *ddpC* | 2.43 | 1.28 | 9.88E-03 | Putative D,D-dipeptide ABC transporter subunit |
| *Z1217* | 2.43 | 1.28 | 0.04 | CP4-44 prophage; RadC-like protein |
| *emrR* | 2.42 | 1.28 | 2.88E-03 | DNA-binding transcriptional regulator |
| *uhpB* | 2.42 | 1.28 | 4.86E-03 | Sensory histidine kinase |
| *lsrB* | 2.41 | 1.27 | 2.56E-06 | Autoinducer-2 ABC transporter periplasmic binding |
| *Z_RS14610* | 2.4 | 1.26 | 7.57E-04 | Hypothetical protein |
| *galM* | 2.39 | 1.26 | 4.40E-06 | Galactose-1-epimerase |

| | | | | |
|---|---|---|---|---|
| *gadB* | 2.38 | 1.25 | 3.75E-06 | Glutamate decarboxylase B |
| *gadA* | 2.38 | 1.25 | 4.21E-06 | Glutamate decarboxylase A |
| *Z0325* | 2.38 | 1.25 | 4.39E-04 | Unknown protein encoded in prophage CP-933I |
| *Z2970* | 2.37 | 1.24 | 4.56E-06 | Putative regulator for prophage CP-933T |
| *cydC* | 2.36 | 1.24 | 5.35E-05 | Glutathione/L-cysteine ABC exporter subunit |
| *guaC* | 2.36 | 1.24 | 2.71E-04 | GMP reductase |
| *pgsA* | 2.34 | 1.23 | 5.45E-05 | CDP-diacylglycerol-glycerol-3-phosphate 3-transferase |
| *pgaA* | 2.32 | 1.21 | 2.42E-05 | Poly-β-1,6-N-acetyl-D-glucosamine export porin |
| *folD* | 2.32 | 1.21 | 1.95E-04 | methylenetetrahydrofolate dehydrogenase |
| *nrdF* | 2.31 | 1.21 | 2.54E-05 | Ribonucleoside-diphosphate reductase 2 subunit β |
| *tadA* | 2.3 | 1.20 | 9.29E-04 | tRNA adenosine34 deaminase |
| *gltK* | 2.28 | 1.19 | 8.10E-05 | Glutamate/aspartate ABC transporter |
| *yeiI* | 2.28 | 1.19 | 1.20E-03 | Putative sugar kinase |
| *Z5882* | 2.28 | 1.19 | 2.28E-03 | Hypothetical protein |
| *yhdX* | 2.27 | 1.18 | 6.75E-04 | Putative ABC transporter membrane subunit |
| *hybG* | 2.27 | 1.18 | 0.04 | Hydrogenase maturation factor |
| *exbD* | 2.26 | 1.18 | 5.73E-05 | Ton complex subunit |
| *rarA* | 2.26 | 1.18 | 1.36E-04 | Replication-associated recombination protein A |
| *bioP* | 2.26 | 1.18 | 2.28E-04 | Biotin transporter |
| *hdeA* | 2.25 | 1.17 | 1.82E-05 | Periplasmic acid stress chaperone |
| *aceE* | 2.25 | 1.17 | 2.70E-05 | Pyruvate dehydrogenase E1 component |
| *alaA* | 2.25 | 1.17 | 3.89E-05 | Glutamate-pyruvate aminotransferase |
| *appC* | 2.25 | 1.17 | 2.00E-03 | Cytochrome *bd*-II subunit 1 |
| *xerC* | 2.24 | 1.16 | 4.86E-04 | Site-specific tyrosine recombinase |
| *yciA* | 2.24 | 1.16 | 4.87E-04 | Acyl-CoA thioesterase |
| *nleF* | 2.23 | 1.16 | 1.18E-04 | T3SS effector |
| *yciY* | 2.23 | 1.16 | 1.51E-04 | Hypothetical protein |
| *nudE* | 2.23 | 1.16 | 1.60E-04 | ADP-sugar diphosphatase |
| *trmJ* | 2.22 | 1.15 | 1.33E-04 | tRNA Cm32/Um32 methyltransferase |
| *recQ* | 2.22 | 1.15 | 1.28E-03 | ATP-dependent DNA helicase |
| *dnaX* | 2.21 | 1.14 | 1.17E-04 | DNA elongation factor III |
| *rapZ* | 2.21 | 1.14 | 3.37E-04 | RNase adaptor protein |
| *Z4273* | 2.21 | 1.14 | 4.20E-03 | P-loop NTPase domain-containing protein |
| *lolD* | 2.21 | 1.14 | 0.02 | Lipoprotein release complex - ATP binding subunit |
| *folE* | 2.2 | 1.14 | 9.11E-05 | GTP cyclohydrolase 1 |
| *dgcP* | 2.2 | 1.14 | 1.33E-04 | Diguanylate cyclase |
| *rhtC* | 2.2 | 1.14 | 5.27E-04 | Threonine export protein |
| *dbpA* | 2.19 | 1.13 | 4.67E-04 | ATP-dependent RNA helicase |
| *yigB* | 2.19 | 1.13 | 4.52E-03 | 5-amino-6- uracil phosphatase |
| *gmd* | 2.19 | 1.13 | 4.84E-03 | GDP-mannose 4,6-dehydratase |
| *xylA* | 2.17 | 1.12 | 7.20E-05 | D-xylose isomerase |
| *fruK* | 2.17 | 1.12 | 8.60E-05 | 1-phosphofructokinase |
| *pbpG* | 2.17 | 1.12 | 4.78E-04 | Peptidoglycan DD-endopeptidase |
| *Z2389* | 2.17 | 1.12 | 8.61E-04 | Putative DNA modification methyltransferase CP-933R |
| *hemP* | 2.17 | 1.12 | 1.92E-03 | Hemin uptake protein |
| *csgC* | 2.17 | 1.12 | 0.05 | Putative curli production protein |
| *lnt* | 2.16 | 1.11 | 1.69E-04 | Apolipoprotein N-acyltransferase |
| *iap* | 2.16 | 1.11 | 3.42E-04 | Alkaline phosphatase isozyme conversion protein |
| *pgaB* | 2.16 | 1.11 | 5.54E-04 | D-glucosamine N-deacetylase |
| *dpaA* | 2.15 | 1.10 | 3.95E-04 | Peptidoglycan meso-diaminopimelicamidase |
| *ysaA* | 2.15 | 1.10 | 1.57E-03 | Putative electron transport protein |

| | | | | |
|---|---|---|---|---|
| *tsaA* | 2.15 | 1.10 | 0.02 | tRNA m6t6A37 methyltransferase |
| *thiH* | 2.15 | 1.10 | 0.04 | 2-iminoacetate synthase |
| *cspA* | 2.14 | 1.10 | 1.15E-04 | Cold shock protein |
| *pdxH* | 2.14 | 1.10 | 8.07E-04 | Pyridoxine/pyridoxamine 5'-phosphate oxidase |
| *lgt* | 2.14 | 1.10 | 2.54E-03 | Phosphatidylglycerol-prolipoprotein transferase |
| *mnmH* | 2.14 | 1.10 | 0.01 | tRNA 2-selenouridine synthase |
| *Z2971* | 2.13 | 1.09 | 7.40E-05 | Unknown protein encoded by prophage CP-933T |
| *Z4317* | 2.13 | 1.09 | 2.31E-04 | Unknown protein encoded by ISEc8 |
| *efeO* | 2.13 | 1.09 | 6.06E-04 | Ferrous iron transport system protein |
| *yfbR* | 2.13 | 1.09 | 2.22E-03 | dCMP phosphohydrolase |
| *yeeO* | 2.12 | 1.08 | 1.55E-04 | Hypothetical protein |
| *rfaF* | 2.12 | 1.08 | 5.60E-04 | ADP-heptose-LPS heptosyltransferase |
| *ycdY* | 2.12 | 1.08 | 1.34E-03 | Chaperone protein |
| *fhuF* | 2.11 | 1.08 | 1.13E-04 | Ferric-siderophore reductase |
| *hutX* | 2.11 | 1.08 | 8.98E-04 | Heme utilization cystosolic carrier protein |
| *Z5881* | 2.11 | 1.08 | 2.22E-03 | Hypothetical protein |
| *ybaP* | 2.1 | 1.07 | 1.59E-04 | TraB family protein |
| *aceF* | 2.1 | 1.07 | 1.59E-04 | Pyruvate dehydrogenase, E2 subunit |
| *Z_RS08590* | 2.1 | 1.07 | 5.35E-03 | Hypothetical protein |
| *yhhJ* | 2.09 | 1.06 | 1.16E-03 | ABC transporter family protein |
| *yeaR* | 2.09 | 1.06 | 1.95E-03 | Hypothetical protein |
| *map* | 2.09 | 1.06 | 4.60E-03 | T3SS effector |
| *tccP* | 2.08 | 1.06 | 5.16E-03 | T3SS effector |
| *entF* | 2.07 | 1.05 | 1.62E-04 | Apo-serine activating enzyme |
| *Z2974* | 2.07 | 1.05 | 1.82E-04 | Unknown protein encoded by prophage CP-933T |
| *fbpC* | 2.07 | 1.05 | 3.11E-03 | CP4-6 prophage; ABC transporter ATP-binding protein |
| *yhiD* | 2.07 | 1.05 | 5.96E-03 | Inner membrane protein |
| *menH* | 2.07 | 1.05 | 0.03 | 2-succinyl-6-hydroxy-2,4-cyclohex-1-carboxylate |
| *csgA* | 2.06 | 1.04 | 1.90E-04 | Curlin, major subunit |
| *Z2991* | 2.06 | 1.04 | 5.88E-04 | Putative tail sheath protein of prophage CP-933T |
| *infA* | 2.06 | 1.04 | 7.02E-04 | Translation initiation factor IF-1 |
| *copD* | 2.06 | 1.04 | 2.88E-03 | CopD family protein |
| *wecC* | 2.06 | 1.04 | 9.28E-03 | UDP-N-acetyl-D-mannosamine dehydrogenase |
| *mltF* | 2.06 | 1.04 | 0.01 | Membrane-bound lytic murein transglycosylase F |
| *Z1341* | 2.05 | 1.04 | 3.66E-04 | Unknown protein prophage CP-933M |
| *Z2979* | 2.05 | 1.04 | 4.96E-04 | Putative stability/partitioning prophage CP-933T |
| *yigL* | 2.05 | 1.04 | 1.84E-03 | Hypothetical protein |
| *escT* | 2.05 | 1.04 | 0.05 | T3SS biogenesis protein |
| *pdhR* | 2.04 | 1.03 | 1.28E-03 | DNA-binding transcriptional dual regulator |
| *mtgA* | 2.04 | 1.03 | 1.33E-03 | Peptidoglycan glycosyltransferase |
| *ispA* | 2.04 | 1.03 | 1.45E-03 | Geranyl diphosphate/farnesyl diphosphate synthas |
| *wcaD* | 2.04 | 1.03 | 0.03 | Colanic acid polymerase |
| *yfaV* | 2.03 | 1.02 | 0.03 | Putative transporter |
| *pstA* | 2.03 | 1.02 | 0.03 | Phosphate ABC transporter membrane subunit |
| *Z5002* | 2.02 | 1.01 | 4.16E-03 | Putative glycoside hydrolase 127 protein |
| *yjcO* | 2.01 | 1.01 | 1.50E-03 | Sel1 repeat-containing protein |
| *ubiI* | 2.01 | 1.01 | 1.95E-03 | 2-octaprenylphenol 6-hydroxylase |
| *Z2387* | 2.01 | 1.01 | 4.48E-03 | Unknown protein encoded within prophage CP-933R |
| *hisP* | 2.01 | 1.01 | 5.51E-03 | Lysine/arginine/ornithine/histidine ABC transporter |
| *btuF* | 2.01 | 1.01 | 0.02 | Vitamin B12 ABC transporter periplasmic binding |
| *sbcB* | 2 | 1.00 | 9.02E-04 | Exodeoxyribonuclease I |

| | | | | |
|---|---|---|---|---|
| *gsk* | 2 | 1.00 | 1.16E-03 | Inosine/guanosine kinase |
| *metK* | 2 | 1.00 | 2.39E-03 | Methionine adenosyltransferase |
| *Z_RS09815* | 2 | 1.00 | 0.04 | DNA-binding protein |
| *serS* | 1.99 | 0.99 | 4.82E-04 | Serine-tRNA ligase |
| *tonB* | 1.98 | 0.99 | 6.38E-04 | Ton complex subunit |
| *Z_RS13965* | 1.98 | 0.99 | 6.69E-04 | Hypothetical protein |
| *minC* | 1.97 | 0.98 | 8.99E-04 | Z-ring positioning protein |
| *ybjX* | 1.97 | 0.98 | 2.50E-03 | DUF535 domain-containing protein |
| *entD* | 1.97 | 0.98 | 9.07E-03 | Phosphopantetheinyl transferase |
| *Z_RS18390* | 1.97 | 0.98 | 0.01 | Hypothetical protein |
| *Z3664* | 1.97 | 0.98 | 0.01 | Putative RNA-guided DNA endonuclease |
| *aspC* | 1.96 | 0.97 | 6.67E-04 | Aspartate aminotransferase |
| *gabT* | 1.96 | 0.97 | 7.71E-04 | 4-aminobutyrate aminotransferase |
| *hemB* | 1.96 | 0.97 | 2.91E-03 | Porphobilinogen synthase |
| *nfi* | 1.96 | 0.97 | 0.02 | Endonuclease V |
| *ppc* | 1.95 | 0.96 | 1.01E-03 | Phosphoenolpyruvate carboxylase |
| *shuT* | 1.95 | 0.96 | 1.42E-03 | Putative periplasmic binding protein |
| *mdtF* | 1.95 | 0.96 | 1.95E-03 | Multidrug efflux pump RND permease |
| *ypfG* | 1.95 | 0.96 | 2.89E-03 | DUF1176 domain-containing protein |
| *potI* | 1.95 | 0.96 | 4.15E-03 | Putrescine ABC transporter membrane subunit |
| *hemG* | 1.95 | 0.96 | 5.41E-03 | Protoporphyrinogen oxidase |
| *rnhB* | 1.95 | 0.96 | 0.04 | RNase HII |
| *yfiR* | 1.93 | 0.95 | 2.17E-03 | DUF4154 domain-containing protein |
| *menC* | 1.93 | 0.95 | 4.45E-03 | *o*-succinylbenzoate synthase |
| *Z2980* | 1.93 | 0.95 | 8.78E-03 | Putative stability/partitioning protein CP-933T |
| *murE* | 1.92 | 0.94 | 2.01E-03 | UDP-N-acetylmuramoyl-L-alanyl-D- ligase |
| *modE* | 1.92 | 0.94 | 9.61E-03 | DNA-binding transcriptional dual regulator |
| *znuC* | 1.91 | 0.93 | 1.23E-03 | Zn2+ ABC transporter ATP binding subunit |
| *yojI* | 1.91 | 0.93 | 3.65E-03 | ABC transporter family protein |
| *Z1533* | 1.91 | 0.93 | 3.71E-03 | Putative oxidoreductase |
| *Z2992* | 1.91 | 0.93 | 4.09E-03 | Putative tail assembly protein of prophage CP-933T |
| *pflC* | 1.91 | 0.93 | 0.04 | Putative pyruvate formate-lyase 2 activating enzyme |
| *tkt_1* | 1.9 | 0.93 | 1.35E-03 | Transketolase 1 |
| *zapC* | 1.9 | 0.93 | 1.52E-03 | Cell division protein |
| *znuB* | 1.9 | 0.93 | 1.92E-03 | Zn2+ ABC transporter membrane subunit |
| *lpxB* | 1.9 | 0.93 | 7.93E-03 | Lipid A disaccharide synthase |
| *nadD* | 1.9 | 0.93 | 0.01 | Nicotinate-nucleotide adenylyltransferase |
| *cutA* | 1.9 | 0.93 | 0.02 | Copper binding protein |
| *aroL* | 1.9 | 0.93 | 0.03 | Shikimate kinase 2 |
| *pyrF* | 1.9 | 0.93 | 0.04 | Orotidine-5'-phosphate decarboxylase |
| *lsrD* | 1.89 | 0.92 | 1.29E-03 | Autoinducer-2 ABC transporter membrane subunit |
| *bioB* | 1.89 | 0.92 | 1.66E-03 | Biotin synthase |
| *Z_RS13970* | 1.89 | 0.92 | 1.82E-03 | Transcriptional regulator |
| *yehS* | 1.89 | 0.92 | 0.01 | DUF1456 domain-containing protein |
| *yeiG* | 1.88 | 0.91 | 2.02E-03 | *S*-formylglutathione hydrolase |
| *tatB* | 1.88 | 0.91 | 2.78E-03 | Sec-independent protein translocase protein |
| *ribD* | 1.88 | 0.91 | 3.12E-03 | Diaminohydroxyphosphoribosylaminopyrimidine |
| *tusD* | 1.88 | 0.91 | 5.00E-03 | Sulphurtransferase complex subunit |
| *lplA* | 1.88 | 0.91 | 0.02 | Outer membrane lipoprotein carrier protein |
| *pstC* | 1.88 | 0.91 | 0.04 | Phosphate ABC transporter membrane subunit |
| *hscA* | 1.87 | 0.90 | 2.18E-03 | Iron-sulfur cluster biosynthesis chaperone |

| | | | | |
|---|---|---|---|---|
| *lapB* | 1.87 | 0.90 | 2.63E-03 | Lipopolysaccharide assembly protein |
| *ratA* | 1.87 | 0.90 | 4.75E-03 | Ribosome association toxin |
| *coaBC* | 1.87 | 0.90 | 7.29E-03 | 4'-phosphopantothenoylcysteine decarboxylase |
| *malQ* | 1.87 | 0.90 | 8.41E-03 | 4-α-glucanotransferase |
| *tyrB* | 1.87 | 0.90 | 9.75E-03 | Tyrosine aminotransferase |
| *pgaC* | 1.87 | 0.90 | 0.01 | Poly-N-acetyl-D-glucosamine synthase subunit |
| *torD* | 1.87 | 0.90 | 0.01 | Trimethylamine-N-oxide reductase-specific chaperone |
| *dedA* | 1.87 | 0.90 | 0.01 | Hypothetical protein |
| *yjfZ* | 1.87 | 0.90 | 0.04 | DUF2686 domain-containing protein |
| *oxyR* | 1.86 | 0.90 | 2.23E-03 | DNA-binding transcriptional regulator |
| *ygfX* | 1.86 | 0.90 | 7.43E-03 | Hypothetical protein |
| *Z2977* | 1.86 | 0.90 | 0.02 | Unknown protein encoded by prophage CP-933T |
| *yrbL* | 1.85 | 0.89 | 3.65E-03 | Protein kinase-like domain-containing protein |
| *rfaL* | 1.85 | 0.89 | 0.01 | *O*-antigen ligase |
| *Z2989* | 1.85 | 0.89 | 0.02 | Unknown protein encoded by prophage CP-933T |
| *apt* | 1.85 | 0.89 | 0.02 | Adenine phosphoribosyltransferase |
| *yfjF* | 1.85 | 0.89 | 0.02 | Putative component of the Rsx system |
| *gabD* | 1.84 | 0.88 | 2.50E-03 | Succinate-semialdehyde dehydrogenase |
| *yagU* | 1.84 | 0.88 | 5.71E-03 | DUF1440 domain-containing inner membrane protein |
| *Z_RS29390* | 1.84 | 0.88 | 6.12E-03 | Hypothetical protein |
| *birA* | 1.84 | 0.88 | 0.01 | Biotin-[acetyl-CoA-carboxylase] ligase |
| *mukE* | 1.84 | 0.88 | 0.01 | Chromosome partitioning protein |
| *aroA* | 1.84 | 0.88 | 0.02 | 3-phosphoshikimate 1-carboxyvinyltransferase |
| *pgaD* | 1.84 | 0.88 | 0.03 | Poly-N-acetyl-D-glucosamine synthase subunit |
| *tusA* | 1.84 | 0.88 | 0.03 | Sulphur transfer protein |
| *lpxA* | 1.83 | 0.87 | 4.75E-03 | Acyl-UDP-N-acetylglucosamine O-acyltransferase |
| *malP* | 1.83 | 0.87 | 7.43E-03 | Maltodextrin phosphorylase |
| *rfaC* | 1.83 | 0.87 | 0.01 | Lipopolysaccharide heptosyltransferase |
| *dam* | 1.83 | 0.87 | 0.01 | DNA adenine methyltransferase |
| *frlD* | 1.83 | 0.87 | 0.03 | Fructoselysine 6-kinase |
| *hscB* | 1.82 | 0.86 | 6.04E-03 | [Fe-S] cluster biosynthesis co-chaperone |
| *yjaH* | 1.82 | 0.86 | 0.01 | DUF1481 domain-containing protein |
| *Z_RS31060* | 1.82 | 0.86 | 0.02 | Transcriptional regulator |
| *Z3269* | 1.82 | 0.86 | 0.02 | Hypothetical protein |
| *yddE* | 1.82 | 0.86 | 0.02 | PF02567 family protein |
| *yacC* | 1.82 | 0.86 | 0.02 | Hypothetical protein |
| *rlmH* | 1.82 | 0.86 | 0.03 | 23S rRNA m3Ψ1915 methyltransferase |
| *udk* | 1.82 | 0.86 | 0.03 | Uridine/cytidine kinase |
| *dnaC* | 1.82 | 0.86 | 0.03 | DNA replication protein |
| *tmaR* | 1.81 | 0.86 | 3.83E-03 | Putative alpha helix protein |
| *rppH* | 1.81 | 0.86 | 4.73E-03 | RNA pyrophosphohydrolase |
| *uvrC* | 1.81 | 0.86 | 6.17E-03 | UvrABC excision nuclease subunit |
| *thrA* | 1.81 | 0.86 | 7.06E-03 | Fused aspartate kinase/homoserine dehydrogenase 1 |
| *mukF* | 1.81 | 0.86 | 8.16E-03 | Chromosome partitioning protein |
| *ybjT* | 1.81 | 0.86 | 0.01 | Putative NAD(P)-binding protein |
| *spoT* | 1.8 | 0.85 | 5.99E-03 | Bifunctional (p)ppGpp synthase/hydrolase |
| *moaC* | 1.8 | 0.85 | 0.01 | Cyclic pyranopterin monophosphate synthase |
| *mdtE* | 1.8 | 0.85 | 0.01 | Multidrug efflux pump membrane fusion protein |
| *rsmB* | 1.8 | 0.85 | 0.01 | 16S rRNA m5C967 methyltransferase |
| *epmB* | 1.8 | 0.85 | 0.02 | Lysine 2,3-aminomutase |
| *espL* | 1.8 | 0.85 | 0.03 | T3SS biogenesis protein |

| | | | | |
|---|---|---|---|---|
| *metB* | 1.8 | 0.85 | 0.03 | *O*-succinylhomoserine(thiol)-lyase |
| *yjjA* | 1.8 | 0.85 | 0.04 | DUF2501 domain-containing protein |
| *glnD* | 1.79 | 0.84 | 6.03E-03 | Protein-PII uridylyltransferase |
| *gss* | 1.79 | 0.84 | 6.13E-03 | glutathionylspermidine amidase/synthetase |
| *recB* | 1.79 | 0.84 | 8.26E-03 | Exodeoxyribonuclease V subunit |
| *Z0957* | 1.79 | 0.84 | 0.02 | Unknown protein encoded by prophage CP-933K |
| *recO* | 1.79 | 0.84 | 0.02 | Recombination mediator protein |
| *mutH* | 1.79 | 0.84 | 0.03 | DNA mismatch repair protein |
| *fre* | 1.78 | 0.83 | 8.29E-03 | NAD(P)H-flavin reductase |
| *dgcN* | 1.78 | 0.83 | 0.01 | Diguanylate cyclase |
| *glyQ* | 1.78 | 0.83 | 0.01 | Glycine-tRNA ligase subunit α |
| *efeB* | 1.78 | 0.83 | 0.01 | Heme-containing peroxidase/deferrochelatase |
| *yeeN* | 1.78 | 0.83 | 0.02 | Putative transcriptional regulator |
| *Z4852* | 1.78 | 0.83 | 0.04 | Putative phospholipid biosynthesis acyltransferase |
| *yfeO* | 1.77 | 0.82 | 8.35E-03 | Ion channel protein |
| *yjjV* | 1.77 | 0.82 | 0.05 | Hypothetical protein |
| *hisS* | 1.76 | 0.82 | 7.77E-03 | Histidine tRNA ligase |
| *ubiH* | 1.76 | 0.82 | 8.29E-03 | 2-octaprenyl-6-methoxyphenol 4-hydroxylase |
| *rapA* | 1.76 | 0.82 | 0.01 | RNAP recycling factor |
| *viaA* | 1.76 | 0.82 | 0.01 | Hypothetical protein |
| *yijE* | 1.76 | 0.82 | 0.02 | Cysteine exporter |
| *ydgC* | 1.76 | 0.82 | 0.02 | Hypothetical protein |
| *yjcB* | 1.76 | 0.82 | 0.02 | Hypothetical protein |
| *shuU* | 1.76 | 0.82 | 0.02 | Heme ABC transporter permease |
| *fumE* | 1.76 | 0.82 | 0.03 | Fumarase E |
| *bcsQ* | 1.75 | 0.81 | 0.02 | Cellulose biosynthesis protein |
| *yibL* | 1.75 | 0.81 | 0.02 | DUF2810 domain-containing protein |
| *fsa* | 1.75 | 0.81 | 0.04 | Fructose-6-phosphate aldolase |
| *mglA* | 1.74 | 0.80 | 7.56E-03 | D-galactose/methyl-galactoside ABC transporter |
| *Z4850* | 1.74 | 0.80 | 0.02 | Putative O-methyltransferase |
| *yceB* | 1.74 | 0.80 | 0.02 | Putative lipid-binding lipoprotein |
| *yshB* | 1.74 | 0.80 | 0.02 | Small membrane protein |
| *yhhS* | 1.74 | 0.80 | 0.02 | Putative transporter |
| *Z2519* | 1.74 | 0.80 | 0.04 | UPF0509 family protein |
| *ppiD* | 1.73 | 0.79 | 9.42E-03 | Periplasmic folding chaperone |
| *Z0309* | 1.73 | 0.79 | 9.70E-03 | Putative cI repressor protein for prophage CP-933H |
| *glyS* | 1.73 | 0.79 | 0.01 | Glycine-tRNA ligase subunit β |
| *topB* | 1.73 | 0.79 | 0.01 | DNA topoisomerase III |
| *atpH* | 1.73 | 0.79 | 0.02 | ATP synthase F1 complex subunit δ |
| *csdA* | 1.73 | 0.79 | 0.02 | Cysteine sulfinate desulfinase |
| *thiQ* | 1.73 | 0.79 | 0.04 | Thiamine ABC transporter ATP binding subunit |
| *Z0308* | 1.73 | 0.79 | 0.05 | Unknown protein from prophage CP-933H |
| *abrB* | 1.73 | 0.79 | 0.05 | Putative regulator |
| *lhgO* | 1.72 | 0.78 | 8.78E-03 | L-2-hydroxyglutarate dehydrogenase |
| *pldB* | 1.72 | 0.78 | 0.01 | Lysophospholipase L2 |
| *trxB* | 1.71 | 0.77 | 0.01 | Thioredoxin reductase |
| *gfcB* | 1.71 | 0.77 | 0.01 | Lipoprotein |
| *ybdZ* | 1.71 | 0.77 | 0.02 | Enterobactin biosynthesis protein |
| *rplK* | 1.7 | 0.77 | 0.01 | 50S ribosomal subunit protein L11 |
| *fruA* | 1.7 | 0.77 | 0.01 | Fructose-specific PTS multiphosphoryl transfer protein |
| *ubiD* | 1.7 | 0.77 | 0.01 | 3-octaprenyl-4-hydroxybenzoate decarboxylase |

| | | | | |
|---|---|---|---|---|
| *glk* | 1.7 | 0.77 | 0.01 | Glucokinase |
| *bisC* | 1.7 | 0.77 | 0.02 | Biotin sulfoxide reductase |
| *tilS* | 1.7 | 0.77 | 0.02 | tRNAIle-lysidine synthetase |
| *Z2975* | 1.7 | 0.77 | 0.02 | Unknown protein encoded by prophage CP-933T |
| *gfcC* | 1.7 | 0.77 | 0.03 | Capsule biosynthesis GfcC family protein |
| *clsB* | 1.7 | 0.77 | 0.03 | Cardiolipin synthase B |
| *ygfI* | 1.7 | 0.77 | 0.04 | Transcriptional regulator |
| *mgtA* | 1.7 | 0.77 | 0.04 | Mg2+ importing P-type ATPase |
| *Z2978* | 1.69 | 0.76 | 0.01 | Putative replication protein for prophage CP-933T |
| *relA* | 1.69 | 0.76 | 0.01 | GDP/GTP pyrophosphokinase |
| *rlmD* | 1.69 | 0.76 | 0.02 | 23S rRNA m5U1939 methyltransferase |
| *alaC* | 1.69 | 0.76 | 0.02 | Glutamate-pyruvate aminotransferase |
| *rng* | 1.69 | 0.76 | 0.02 | RNase G |
| *dcm* | 1.69 | 0.76 | 0.03 | DNA-cytosine methyltransferase |
| *wzxE* | 1.69 | 0.76 | 0.03 | Lipid IIIECA flippase |
| *bfd* | 1.69 | 0.76 | 0.03 | Bacterioferritin-associated ferredoxin |
| *mdoC* | 1.69 | 0.76 | 0.04 | Osmoregulated periplasmic glucans biosynthesis |
| *hrpB* | 1.68 | 0.75 | 0.02 | RNA-dependent NTPase |
| *Z3931* | 1.68 | 0.75 | 0.04 | Unknown protein encoded by prophage CP-933Y |
| *glrR* | 1.68 | 0.75 | 0.05 | DNA-binding transcriptional activator |
| *iscR* | 1.67 | 0.74 | 0.01 | DNA-binding transcriptional dual regulator |
| *ptsP* | 1.67 | 0.74 | 0.03 | Phosphoenolpyruvate-protein phosphotransferase |
| *ygaZ* | 1.67 | 0.74 | 0.03 | L-valine exporter |
| *Z2990* | 1.67 | 0.74 | 0.04 | Putative tail fiber component of prophage CP-933T |
| *hisM* | 1.67 | 0.74 | 0.04 | Lysine/arginine/ornithine/histidine ABC transporter |
| *Z_RS32395* | 1.66 | 0.73 | 0.01 | Hypothetical protein |
| *ycbX* | 1.66 | 0.73 | 0.02 | 6-N-hydroxylaminopurine resistance protein |
| *ddlA* | 1.66 | 0.73 | 0.02 | D-alanine-D-alanine ligase A |
| *can* | 1.66 | 0.73 | 0.02 | Carbonic anhydrase 2 |
| *exbB* | 1.66 | 0.73 | 0.02 | Ton complex subunit |
| *acrA* | 1.66 | 0.73 | 0.03 | Multidrug efflux pump membrane fusion lipoprotein |
| *yqiA* | 1.66 | 0.73 | 0.03 | Esterase |
| *polB* | 1.65 | 0.72 | 0.02 | DNA polymerase II |
| *rpsF* | 1.65 | 0.72 | 0.02 | 30S ribosomal subunit protein S6 |
| *yjbH* | 1.65 | 0.72 | 0.03 | Putative lipoprotein |
| *murF* | 1.65 | 0.72 | 0.03 | Dalanyl-D-alanine-adding enzyme |
| *rlmB* | 1.65 | 0.72 | 0.03 | 23S rRNA 2'-O-ribose G2251 methyltransferase |
| *dapF* | 1.65 | 0.72 | 0.03 | Diaminopimelate epimerase |
| *mnmC* | 1.65 | 0.72 | 0.03 | 5-aminomethyl-2-thiouridylate methyltransferase |
| *lolA* | 1.65 | 0.72 | 0.03 | Outer membrane lipoprotein carrier protein |
| *mnmE* | 1.65 | 0.72 | 0.04 | 5-carboxymethylaminomethyluridine-tRNA synthase |
| *Z4385* | 1.65 | 0.72 | 0.04 | Putative ATP-binding protein of ABC transporter family |
| *pxpA* | 1.65 | 0.72 | 0.04 | 5-oxoprolinase component A |
| *rplD* | 1.64 | 0.71 | 0.02 | 50S ribosomal subunit protein L4 |
| *fbaA* | 1.64 | 0.71 | 0.02 | Fructose-bisphosphate aldolase class II |
| *yfgM* | 1.64 | 0.71 | 0.03 | Ancillary SecYEG translocon subunit |
| *hemE* | 1.64 | 0.71 | 0.03 | Uroporphyrinogen decarboxylase |
| *mdtH* | 1.64 | 0.71 | 0.04 | Hypothetical protein |
| *priB* | 1.64 | 0.71 | 0.04 | Primosomal replication protein N |
| *mukB* | 1.63 | 0.70 | 0.02 | Chromosome partitioning protein |
| *ubiB* | 1.63 | 0.70 | 0.03 | Ubiquinone biosynthesis protein |

| | | | | |
|---|---|---|---|---|
| *Z2985* | 1.63 | 0.70 | 0.04 | Putative tail fiber protein of prophage CP-933T |
| *Z1519* | 1.63 | 0.70 | 0.04 | Hypothetical protein |
| *dnaA* | 1.62 | 0.70 | 0.02 | Chromosomal replication initiator protein |
| *lsrC* | 1.62 | 0.70 | 0.02 | Autoinducer-2 ABC transporter membrane subunit |
| *ribF* | 1.62 | 0.70 | 0.03 | Bifunctional riboflavin kinase |
| *gpmM* | 1.62 | 0.70 | 0.03 | phosphoglycerate mutase |
| *dapA* | 1.62 | 0.70 | 0.03 | 4-hydroxy-tetrahydrodipicolinate synthase |
| *yqhD* | 1.61 | 0.69 | 0.03 | NADPH-dependent aldehyde reductase |
| *pepD* | 1.61 | 0.69 | 0.03 | Peptidase D |
| *rplA* | 1.61 | 0.69 | 0.03 | 50S ribosomal subunit protein L1 |
| *zinT* | 1.61 | 0.69 | 0.03 | Metal-binding protein |
| *espG* | 1.61 | 0.69 | 0.04 | T3SS effector |
| *rbbA* | 1.61 | 0.69 | 0.04 | Ribosome-associated ATPase |
| *iscS* | 1.6 | 0.68 | 0.03 | Cysteine desulfurase |
| *cyaA* | 1.6 | 0.68 | 0.03 | Adenylate cyclase |
| *nanR* | 1.6 | 0.68 | 0.04 | Transcriptional regulator |
| *galR* | 1.6 | 0.68 | 0.04 | DNA-binding transcriptional dual regulator |
| *cpxA* | 1.59 | 0.67 | 0.03 | Sensor histidine kinase |
| *iscU* | 1.59 | 0.67 | 0.04 | Scaffold protein for iron-sulfur cluster assembly |
| *cvrA* | 1.58 | 0.66 | 0.04 | Potassium/proton antiporter |
| *rpsK* | 1.58 | 0.66 | 0.04 | 30S ribosomal subunit protein S11 |
| *cydA* | 1.58 | 0.66 | 0.04 | Cytochrome bd-I subunit 1 |
| *maeA* | 1.58 | 0.66 | 0.05 | Malate dehydrogenase |
| *pdxA* | 1.58 | 0.66 | 0.05 | 4-hydroxythreonine-4-phosphate dehydrogenase |
| *rpsS* | 1.58 | 0.66 | 0.05 | 30S ribosomal subunit protein S19 |
| *sthA* | 1.57 | 0.65 | 0.03 | Soluble pyridine nucleotide transhydrogenase |
| *rplC* | 1.57 | 0.65 | 0.04 | 50S ribosomal subunit protein L3 |
| *nuoC* | 1.57 | 0.65 | 0.04 | NADH:quinone oxidoreductase subunit |
| *hpf* | 1.57 | 0.65 | 0.04 | Ribosome hibernation-promoting factor |
| *hrpA* | 1.57 | 0.65 | 0.04 | ATP-dependent 3'→5' RNA helicase |
| *prfB* | 1.57 | 0.65 | 0.05 | Peptide chain release factor RF2 |
| *Z2987* | 1.57 | 0.65 | 0.05 | Putative tail fiber component of prophage CP-933T |
| *rplB* | 1.56 | 0.64 | 0.04 | 50S ribosomal subunit protein L2 |
| *nrdE* | 1.56 | 0.64 | 0.04 | Ribonucleoside-diphosphate reductase 2 subunit α |
| *mpl* | 1.56 | 0.64 | 0.05 | N-acetylmuramate-L-alanyl-γ-D-glutamyl-meso ligase |
| *minD* | 1.55 | 0.63 | 0.05 | Z-ring positioning protein |
| *dctA* | 1.54 | 0.62 | 0.04 | C4 dicarboxylate/orotate:H+ symporter |
| *bamA* | 1.54 | 0.62 | 0.05 | Outer membrane protein assembly factor |
| ***Downregulated*** | | | | |
| *garP* | -14.7 | -3.88 | 3.98E-32 | Galactarate/D-glucarate transporter |
| *mgtS* | -12.91 | -3.69 | 1.95E-43 | Hypothetical protein |
| *yjfN* | -12.2 | -3.61 | 0 | Protease inhibitor |
| *fadD* | -11.32 | -3.50 | 0 | Long-chain-fatty-acid—CoA ligase |
| *mntS* | -11 | -3.46 | 0 | Manganase accumulation protein |
| *idlP* | -10.43 | -3.38 | 0.04 | Leader peptide |
| *bsmA* | -9.67 | -3.27 | 0 | Biofilm peroxide resistance protein |
| *murQ* | -8.8 | -3.14 | 5.02E-29 | N-acetylmuramic acid 6-phosphate etherase |
| *putA* | -8.39 | -3.07 | 9.24E-30 | Proline dehydrogenase, P5C dehydrogenase |
| *Z5589* | -8.06 | -3.01 | 3.10E-30 | 23S ribosomal RNA |
| *Z_RS12265* | -7.53 | -2.91 | 6.29E-11 | Hypothetical protein |
| *Z4637* | -7.45 | -2.90 | 6.05E-24 | 23S ribosomal RNA |

| | | | | |
|---|---|---|---|---|
| *garL* | -7.37 | -2.88 | 3.85E-16 | α-dehydro-β-deoxy-D-glucarate aldolase |
| *Z3875* | -6.83 | -2.77 | 1.21E-19 | 23S ribosomal RNA |
| *mcbA* | -6.78 | -2.76 | 9.52E-25 | DUF1471 family periplasmic protein |
| *fadE* | -6.73 | -2.75 | 0 | acyl-CoA dehydrogenase |
| *gtdA* | -6.49 | -2.70 | 3.18E-21 | Putative 1,2-dioxygenase |
| *pspG* | -6.47 | -2.69 | 0 | Phage shock protein G |
| *fadL* | -6.36 | -2.67 | 0 | Long-chain fatty acid outer membrane channel |
| *Z5259* | -6.34 | -2.66 | 4.07E-04 | 16S ribosomal RNA |
| *ybeL* | -6.28 | -2.65 | 0 | DUF1451 domain-containing protein |
| *ynfM* | -6.22 | -2.64 | 0 | Putative transport protein |
| *cysD* | -6.19 | -2.63 | 9.18E-21 | Sulphate adenylyltransferase subunit 2 |
| *Z3392* | -6.07 | -2.60 | 7.00E-20 | Putative isomerase-decarboxylase |
| *ivbL* | -6.01 | -2.59 | 0.04 | Leader peptide |
| *Z5379* | -6.01 | -2.59 | 3.85E-15 | 23S ribosomal RNA |
| *Z0219* | -5.96 | -2.58 | 0 | 23S ribosomal RNA |
| *ugpB* | -5.94 | -2.57 | 0 | sn-glycerol 3-phosphate ABC transporter periplasmic |
| *bolA* | -5.93 | -2.57 | 0 | Putative regulator of murein genes |
| *grcA* | -5.85 | -2.55 | 0 | Stress-induced alternate pyruvate formate-lyase |
| *ydjF* | -5.84 | -2.55 | 4.91E-21 | Putative DNA-binding transcriptional regulator |
| *fadI* | -5.83 | -2.54 | 0 | 3-ketoacyl-CoA thiolase |
| *yodC* | -5.77 | -2.53 | 3.73E-21 | Hypothetical protein |
| *ydcJ* | -5.68 | -2.51 | 0 | VOC family protein |
| *maiA* | -5.64 | -2.50 | 2.48E-17 | Putative glutathione-S-transferase |
| *yejG* | -5.57 | -2.48 | 0 | Hypothetical protein |
| *Z5534* | -5.56 | -2.48 | 4.20E-20 | 23S ribosomal RNA |
| *cutC* | -5.54 | -2.47 | 0 | Copper homeostasis protein |
| *yohP* | -5.53 | -2.47 | 2.88E-03 | Small membrane protein |
| *ychH* | -5.29 | -2.40 | 0 | Stress-induced protein |
| *nrdD* | -5.27 | -2.40 | 0 | Anaerobic ribonucleoside-triphosphate reductase |
| *ydfA* | -5.24 | -2.39 | 0.05 | Putative sulfatase |
| *murP* | -5.22 | -2.38 | 1.01E-17 | N-acetylmuramic acid-specific PTS enzyme |
| *actP* | -5.09 | -2.35 | 2.52E-18 | Acetate/glycolate:cation symporter |
| *cysP* | -5.05 | -2.34 | 4.10E-18 | Thiosulfate/sulphate ABC transporter periplasmic |
| *dhaK* | -5.05 | -2.34 | 0 | Dihydroxyacetone kinase subunit K |
| *acs* | -5.03 | -2.33 | 0 | Acetyl-CoA synthetase |
| *Z4018* | -4.95 | -2.31 | 0 | Putative flavodoxin |
| *fadB* | -4.94 | -2.30 | 0 | L-3-hydroxyacyl-CoA dehydrogenase |
| *fadH* | -4.89 | -2.29 | 1.50E-17 | 2,4-dienoyl-CoA reductase |
| *yfeK* | -4.86 | -2.28 | 1.29E-11 | Hypothetical protein |
| *pbp4b* | -4.77 | -2.25 | 4.72E-16 | Penicillin binding protein 4B |
| *pspC* | -4.76 | -2.25 | 0 | Phage shock protein C |
| *pspB* | -4.74 | -2.24 | 0 | Phage shock protein B |
| *pspE* | -4.74 | -2.24 | 0 | Phage shock protein E |
| *dkgA* | -4.68 | -2.23 | 0 | 2,5-didehydrogluconate reductase |
| *YibT* | -4.67 | -2.22 | 0 | Putative RNAse adapter protein |
| *sseA* | -4.65 | -2.22 | 0 | 3-mercaptopyruvate sulfurtransferase |
| *ldtD* | -4.62 | -2.21 | 0 | L,D-transpeptidase |
| *adrA* | -4.56 | -2.19 | 1.44E-15 | Diguanylate cyclase |
| *yacL* | -4.55 | -2.19 | 0 | Hypothetical protein |
| *yjgR* | -4.54 | -2.18 | 0 | DUF853 domain-containing protein |
| *Z4090* | -4.47 | -2.16 | 0.04 | Hypothetical protein |

| | | | | |
|---|---|---|---|---|
| *ybhQ* | -4.46 | -2.16 | 0 | Inner membrane protein |
| *aceA* | -4.41 | -2.14 | 1.04E-14 | Isocitrate lyase |
| *dhaL* | -4.4 | -2.14 | 4.05E-15 | Dihydroxyacetone kinase subunit L |
| *pspA* | -4.38 | -2.13 | 0 | Phage shock protein A |
| *pspD* | -4.31 | -2.11 | 0 | Phage shock protein D |
| *ybiI* | -4.29 | -2.10 | 0 | Zinc finger domain-containing protein |
| *norW* | -4.16 | -2.06 | 2.04E-14 | NADH:flavorubredoxin reductase |
| *yeaY* | -4.13 | -2.05 | 9.95E-14 | Slp family lipoprotein |
| *puuD* | -4.06 | -2.02 | 9.08E-12 | γ-glutamyl-γ-aminobutyrate hydrolase |
| *Z3394* | -4.06 | -2.02 | 3.61E-13 | Putative transporter |
| *ygjR* | -4.05 | -2.02 | 2.39E-14 | Putative oxidoreductase |
| *osmE* | -4 | -2.00 | 3.52E-14 | Osmotically inducible lipoprotein |
| *melR* | -3.99 | -2.00 | 3.52E-14 | DNA-binding transcriptional dual regulator |
| *Z4645* | -3.98 | -1.99 | 4.80E-03 | 16S ribosomal RNA |
| *aceB* | -3.92 | -1.97 | 1.52E-12 | Malate synthase |
| *prpR* | -3.88 | -1.96 | 2.20E-13 | Regulator for prp operon |
| *dhaM* | -3.86 | -1.95 | 2.38E-13 | Dihydroxyacetone kinase subunit M |
| *sbmC* | -3.86 | -1.95 | 1.79E-13 | Putrescine ABC exporter membrane subunit |
| *tcyJ* | -3.86 | -1.95 | 2.11E-13 | Cysteine ABC transporter substrate-binding protein |
| *Z3882* | -3.86 | -1.95 | 5.37E-10 | 16S ribosomal RNA |
| *ycgB* | -3.79 | -1.92 | 3.21E-13 | PF04293 family protein |
| *hmpA* | -3.76 | -1.91 | 6.15E-13 | Nitric oxide dioxygenase |
| *bssR* | -3.75 | -1.91 | 4.38E-13 | Regulator of biofilm formation |
| *chpS* | -3.75 | -1.91 | 2.14E-11 | ChpS antitoxin, ChpB-ChpS toxin-antitoxin system |
| *cstA* | -3.74 | -1.90 | 4.86E-13 | Carbon starvation protein |
| *ynhF* | -3.72 | -1.90 | 1.57E-11 | Cytochrome bd-I accessory subunit |
| *Z4353* | -3.68 | -1.88 | 1.93E-12 | Putative enzyme |
| *frdD* | -3.67 | -1.88 | 3.77E-12 | Fumarate reductase membrane protein |
| *yqaE* | -3.66 | -1.87 | 1.46E-12 | Pmp3 family protein |
| *ldtE* | -3.65 | -1.87 | 1.48E-12 | L,D-transpeptidase |
| *fadA* | -3.63 | -1.86 | 3.51E-12 | Thiolase I |
| *yobB* | -3.62 | -1.86 | 4.27E-12 | Putative carbon-nitrogen hydrolase family protei |
| *ybeQ* | -3.6 | -1.85 | 6.88E-12 | Sel1 repeat-containing protein |
| *fadJ* | -3.56 | -1.83 | 3.96E-12 | Multifunctional 3-hydroxyacyl-CoA dehydrogenase |
| *allR* | -3.5 | -1.81 | 8.45E-12 | DNA-binding transcriptional repressor |
| *yodD* | -3.5 | -1.81 | 7.17E-12 | Stress-induced protein |
| *dinJ* | -3.49 | -1.80 | 1.49E-11 | Antitoxin/DNA-binding transcriptional repressor |
| *folA* | -3.49 | -1.80 | 7.04E-11 | Dihydrofolate reductase |
| *ilvN* | -3.48 | -1.80 | 6.90E-07 | Acetohydroxy acid synthase I subunit |
| *mhpR* | -3.48 | -1.80 | 3.40E-11 | DNA-binding transcriptional activator |
| *yfdY* | -3.48 | -1.80 | 1.72E-11 | DUF2545 domain-containing protein |
| *htpX* | -3.47 | -1.79 | 1.18E-11 | Protease |
| *ynfD* | -3.45 | -1.79 | 1.48E-11 | DUF1161 domain-containing protein |
| *putP* | -3.44 | -1.78 | 3.03E-11 | Major sodium/proline symporter |
| *garR* | -3.43 | -1.78 | 1.45E-09 | Tartronate semialdehyde reductase |
| *Z_RS32125* | -3.41 | -1.77 | 2.11E-11 | Hypothetical protein |
| *ataR* | -3.37 | -1.75 | 4.00E-11 | Hypothetical protein |
| *cysN* | -3.34 | -1.74 | 2.55E-10 | Sulphate adenylyltransferase subunit 1 |
| *kdpD* | -3.33 | -1.74 | 8.01E-11 | Sensor histidine kinase |
| *yajO* | -3.33 | -1.74 | 7.02E-11 | 1-deoxyxylulose-5-phosphate synthase |
| *yfcH* | -3.33 | -1.74 | 7.32E-11 | Epimerase family protein |

| | | | | |
|---|---|---|---|---|
| *Z5684* | -3.33 | -1.74 | 1.35E-10 | Putative transcriptional regulator |
| *yceH* | -3.28 | -1.71 | 2.62E-10 | DUF480 domain-containing protein |
| *ytfH* | -3.28 | -1.71 | 1.41E-09 | Hypothetical protein |
| *yhaH* | -3.26 | -1.70 | 1.30E-10 | Putative inner membrane protein |
| *aldB* | -3.25 | -1.70 | 1.65E-10 | Aldehyde dehydrogenase B |
| *yhhA* | -3.25 | -1.70 | 1.55E-10 | DUF2756 domain-containing protein |
| *ygaC* | -3.22 | -1.69 | 8.31E-10 | DUF2002 domain-containing protein |
| *yidQ* | -3.22 | -1.69 | 1.22E-09 | Hypothetical protein |
| *dnaK* | -3.21 | -1.68 | 2.02E-10 | Chaperone Hsp70 |
| *yiaG* | -3.21 | -1.68 | 2.67E-10 | Putative DNA-binding transcriptional regulator |
| *csiE* | -3.19 | -1.67 | 2.58E-10 | Stationary phase-inducible protein |
| *yhfG* | -3.19 | -1.67 | 9.54E-09 | DUF2559 domain-containing protein |
| *yjiJ* | -3.17 | -1.66 | 1.06E-09 | Putative transporter |
| *Z1059* | -3.16 | -1.66 | 7.41E-10 | Hypothetical protein |
| *ygaV* | -3.15 | -1.66 | 2.13E-08 | Putative DNA-binding transcriptional regulator |
| *ytfJ* | -3.14 | -1.65 | 1.42E-09 | PF09695 family protein |
| *frdC* | -3.13 | -1.65 | 2.57E-09 | Fumarate reductase membrane protein |
| *qmcA* | -3.13 | -1.65 | 5.97E-10 | PHB domain-containing protein |
| *yaiA* | -3.11 | -1.64 | 2.12E-09 | Hypothetical protein |
| *ihfA* | -3.1 | -1.63 | 6.87E-10 | Integration host factor subunit α |
| *yraR* | -3.1 | -1.63 | 9.16E-10 | Putative nucleoside-diphosphate-sugar epimerase |
| *cpxP* | -3.06 | -1.61 | 1.27E-09 | Periplasmic protein |
| *yohC* | -3.06 | -1.61 | 1.22E-09 | Putative inner membrane protein |
| *folC* | -3.05 | -1.61 | 2.44E-09 | Bifunctional folylpolyglutamate synthetase / dihydrofolate synthetase |
| *dauA* | -3.04 | -1.60 | 1.85E-09 | Aerobic C4-dicarboxylate transporte |
| *agp* | -3.03 | -1.60 | 1.60E-09 | Glucose-1-phosphatase |
| *cysJ* | -3.03 | -1.60 | 8.38E-08 | Sulphite reductase, flavoprotein subunit |
| *ssb1* | -3.03 | -1.60 | 2.39E-09 | Single-stranded DNA-binding protein |
| *yqjG* | -3.03 | -1.60 | 3.26E-09 | Glutathionyl-hydroquinone reductase |
| *srlR* | -3.02 | -1.59 | 1.37E-08 | DNA-binding transcriptional repressor |
| *ssrA* | -3.02 | -1.59 | 1.75E-09 | Transfer-messenger RNA |
| *yeaG* | -3 | -1.58 | 2.40E-09 | Protein kinase |
| *groS* | -2.99 | -1.58 | 4.59E-09 | Co-chaperonin GroES |
| *Z1060* | -2.99 | -1.58 | 3.41E-09 | Hypothetical protein |
| *ahr* | -2.98 | -1.58 | 1.41E-08 | NADPH-dependent aldehyde reductase |
| *sodC_2* | -2.98 | -1.58 | 3.46E-09 | Superoxide dismutase family protein |
| *btsT* | -2.97 | -1.57 | 9.22E-09 | Pyruvate:H+ symporter |
| *uspE* | -2.96 | -1.57 | 4.05E-09 | Universal stress protein E |
| *tomB* | -2.95 | -1.56 | 5.02E-09 | Hha toxicity modulator |
| *puuA* | -2.93 | -1.55 | 4.04E-08 | Putative glutamine synthetase |
| *uspB* | -2.93 | -1.55 | 5.59E-09 | Universal stress protein B |
| *Z3624* | -2.93 | -1.55 | 2.42E-07 | D-fructokinase |
| *ribB* | -2.92 | -1.55 | 7.61E-09 | 3,4-dihydroxy-2-butanone-4-phosphate synthase |
| *trxC* | -2.92 | -1.55 | 1.40E-08 | Reduced thioredoxin 2 |
| *hcp* | -2.9 | -1.54 | 7.60E-06 | S-nitrosylase |
| *ybcK* | -2.89 | -1.53 | 4.07E-07 | DLP12 prophage; putative recombinase |
| *ydhL* | -2.89 | -1.53 | 1.58E-08 | DUF1289 domain-containing protein |
| *yihS* | -2.89 | -1.53 | 8.55E-06 | Sulfoquinovose isomerase |
| *dsrB* | -2.88 | -1.53 | 8.17E-07 | Hypothetical protein |
| *ydjL* | -2.88 | -1.53 | 4.15E-08 | Putative zinc-binding dehydrogenas |

| | | | | |
|---|---|---|---|---|
| *Z4789* | -2.87 | -1.52 | 4.59E-07 | Hypothetical protein |
| *galF* | -2.86 | -1.52 | 1.74E-08 | UTP:glucose-1-phosphate uridylyltransferase |
| *nrfD* | -2.86 | -1.52 | 0.03 | Formate-dependent nitrate reductase complex |
| *dosC* | -2.85 | -1.51 | 1.59E-08 | DosC-DosP complex |
| *wrbA* | -2.85 | -1.51 | 1.45E-08 | NAD(P)H:quinone oxidoreductase |
| *ascF* | -2.84 | -1.51 | 7.35E-08 | β-glucoside specific PTS enzyme IIBC component |
| *ytfK* | -2.84 | -1.51 | 1.73E-08 | Hypothetical protein |
| *Z4340* | -2.84 | -1.51 | 4.49E-07 | Unknown protein encoded by ISEc8 |
| *Z5087* | -2.84 | -1.51 | 2.69E-08 | Putative integrase, prophage 933L/LEE |
| *def* | -2.83 | -1.50 | 1.99E-08 | Peptide deformylase |
| *mokC* | -2.83 | -1.50 | 2.15E-08 | Regulatory protein |
| *ypfN* | -2.82 | -1.50 | 2.35E-08 | PF13980 family protein |
| *Z2148* | -2.82 | -1.50 | 6.62E-04 | Unknown protein encoded within prophage CP-933O |
| *rimJ* | -2.81 | -1.49 | 4.73E-08 | Ribosomal-protein-S5-alanine N-acetyltransferase |
| *ygaM* | -2.81 | -1.49 | 2.75E-08 | DUF883 domain-containing protein |
| *glpD* | -2.8 | -1.49 | 2.59E-08 | Aerobic glycerol 3-phosphate dehydrogenase |
| *bssS* | -2.79 | -1.48 | 2.77E-08 | Regulator of biofilm formation |
| *cspD* | -2.79 | -1.48 | 2.86E-08 | DNA replication inhibitor |
| *pgl* | -2.79 | -1.48 | 5.11E-08 | 6-phosphogluconolactonase |
| *yccJ* | -2.79 | -1.48 | 3.21E-08 | PF13993 family protein |
| *ybhL* | -2.78 | -1.48 | 3.38E-08 | Bax1-I family protein |
| *Z5009* | -2.78 | -1.48 | 4.09E-08 | Hypothetical protein |
| *ymgE* | -2.77 | -1.47 | 9.14E-08 | PF04226 family protein |
| *csrA* | -2.75 | -1.46 | 4.22E-08 | Carbon storage regulator |
| *ydcK* | -2.75 | -1.46 | 1.32E-07 | Putative acyltransferase |
| *cysI* | -2.74 | -1.45 | 1.60E-06 | Sulphite reductase, hemoprotein subunit |
| *pdxI* | -2.74 | -1.45 | 7.93E-08 | Pyridoxal reductase |
| *ybaV* | -2.74 | -1.45 | 1.32E-05 | Hypothetical protein |
| *htpG* | -2.72 | -1.44 | 7.24E-08 | Chaperone protein |
| *Z0509* | -2.71 | -1.44 | 1.85E-07 | Hypothetical protein |
| *Z0974* | -2.71 | -1.44 | 2.46E-03 | Putative tail component of prophage CP-933K |
| *csqR* | -2.7 | -1.43 | 1.21E-06 | DNA-binding transcriptional dual regulator |
| *Z3620* | -2.69 | -1.43 | 9.71E-08 | Hypothetical protein |
| *nudL* | -2.68 | -1.42 | 1.13E-05 | Putative NUDIX hydrolase |
| *yobF* | -2.68 | -1.42 | 1.14E-07 | DUF2527 domain-containing protein |
| *chpB* | -2.66 | -1.41 | 4.49E-07 | Endoribonuclease toxin |
| *pat* | -2.66 | -1.41 | 1.30E-07 | Protein lysine acetyltransferase |
| *cysT* | -2.64 | -1.40 | 1.69E-06 | tRNA-Cys |
| *hokB* | -2.64 | -1.40 | 2.17E-07 | hokB |
| *yjfY* | -2.64 | -1.40 | 6.47E-07 | DUF1471 domain-containing protein |
| *yniA* | -2.63 | -1.40 | 2.26E-07 | Putative kinase |
| *kdgR* | -2.62 | -1.39 | 2.78E-07 | DNA-binding transcriptional repressor |
| *dacC* | -2.61 | -1.38 | 3.02E-07 | D-alanyl-D-alanine carboxypeptidase |
| *iraD* | -2.61 | -1.38 | 1.39E-03 | Anti-adaptor protein |
| *yfiL* | -2.61 | -1.38 | 2.66E-06 | Hypothetical protein |
| *yjgA* | -2.61 | -1.38 | 3.03E-07 | Putative ribosome biogenesis factor |
| *acnA* | -2.6 | -1.38 | 3.08E-07 | Aconitate hydrase 1 |
| *hokD_3* | -2.6 | -1.38 | 2.91E-07 | Type I toxin-antitoxin system toxin |
| *ydcL* | -2.6 | -1.38 | 2.94E-07 | DUF3313 domain-containing lipoprotein |
| *yhhT* | -2.6 | -1.38 | 7.59E-07 | Hypothetical protein |
| *ytjA* | -2.6 | -1.38 | 4.05E-07 | Hypothetical protein |

| | | | | |
|---|---|---|---|---|
| *rnpB* | -2.59 | -1.37 | 2.97E-07 | RNase P catalytic RNA component |
| *yoaC* | -2.59 | -1.37 | 1.00E-05 | Hypothetical protein |
| *Z5086* | -2.59 | -1.37 | 0.03 | tRNA |
| *zntB* | -2.58 | -1.37 | 6.01E-07 | Zn2+:H+ symporte |
| *treA* | -2.57 | -1.36 | 9.17E-07 | Alpha,alpha-trehalase |
| *ybdK* | -2.57 | -1.36 | 4.80E-07 | Putative glutamate cysteine ligase 2 |
| *Z1924* | -2.56 | -1.36 | 1.05E-06 | Stress-induced acidophilic repeat motifs protein |
| *Z_RS33015* | -2.54 | -1.34 | 6.04E-07 | Hypothetical protein |
| *glgS* | -2.53 | -1.34 | 7.27E-07 | Surface composition regulator |
| *yniB* | -2.53 | -1.34 | 7.06E-07 | Uncharacterised protein |
| *caiF* | -2.52 | -1.33 | 3.30E-05 | Transcriptional regulator |
| *panE* | -2.52 | -1.33 | 9.30E-07 | 2-dehydropantoate 2-reductase |
| *psiF* | -2.52 | -1.33 | 1.57E-06 | Phosphate starvation-inducible protein |
| *rhaS* | -2.52 | -1.33 | 5.05E-05 | Rranscriptional activator |
| *ygfZ* | -2.52 | -1.33 | 1.03E-06 | Folate-binding protein |
| *yibI* | -2.52 | -1.33 | 8.43E-06 | DUF3302 domain-containing protein |
| *Z5150* | -2.52 | -1.33 | 1.91E-06 | Hypothetical protein |
| *sdaA* | -2.51 | -1.33 | 8.40E-07 | L-serine deaminase |
| *ubiF* | -2.51 | -1.33 | 1.03E-06 | 3-demethoxyubiquinol 3-hydroxylase |
| *yeaX* | -2.51 | -1.33 | 9.93E-06 | Carnitine monooxygenase subunit |
| *yihU* | -2.51 | -1.33 | 3.95E-03 | 3-sulfolactaldehyde reductase |
| *yhgE* | -2.5 | -1.32 | 4.24E-06 | Putative transporter |
| *gatY* | -2.48 | -1.31 | 0.03 | Tagatose-1,6-bisphosphate aldolase 2 |
| *groL* | -2.48 | -1.31 | 1.17E-06 | Chaperonin GroEL |
| *uhpT* | -2.48 | -1.31 | 2.10E-05 | Hexose-6-phosphate:phosphate antiporter |
| *yccU* | -2.48 | -1.31 | 1.87E-06 | Putative HspQ acetyl donor |
| *ydcS* | -2.48 | -1.31 | 1.75E-06 | Putative ABC transporter periplasmic binding protein |
| *ygiW* | -2.48 | -1.31 | 1.31E-06 | BOF family protein |
| *xylG* | -2.47 | -1.30 | 2.56E-06 | Xylose ABC transporter ATP binding subunit |
| *Z0771* | -2.47 | -1.30 | 1.78E-06 | Hypothetical protein |
| *nimT* | -2.46 | -1.30 | 1.34E-05 | 2-nitroimidazole exporter |
| *yeaH* | -2.46 | -1.30 | 1.51E-06 | DUF444 domain-containing protein |
| *yicJ* | -2.46 | -1.30 | 8.84E-06 | Putative permease |
| *rbsD* | -2.45 | -1.29 | 1.85E-06 | D-ribose pyranase |
| *asnC* | -2.44 | -1.29 | 1.37E-04 | DNA-binding transcriptional dual regulator |
| *osmB* | -2.43 | -1.28 | 2.31E-06 | Osmotically inducible lipoprotein |
| *uspG* | -2.42 | -1.28 | 2.67E-06 | Universal stress protein G |
| *hspQ* | -2.41 | -1.27 | 2.56E-06 | Putative HspQ acetyl donor |
| *kdpE* | -2.41 | -1.27 | 6.03E-06 | DNA-binding transcriptional activator |
| *rffT* | -2.41 | -1.27 | 1.57E-05 | N-acetylfucosaminyltransferase |
| *Z_RS14675* | -2.41 | -1.27 | 3.91E-06 | LexA family transcriptional regulator |
| *cysH* | -2.4 | -1.26 | 1.05E-04 | phosphoadenosine phosphosulphate reductase |
| *agaV* | -2.39 | -1.26 | 0.03 | N-acetyl-D-galactosamine specific PTS |
| *robA* | -2.39 | -1.26 | 3.82E-06 | DNA-binding transcriptional dual regulator |
| *yphG* | -2.39 | -1.26 | 1.62E-05 | DUF5107 domain-containing protein |
| *cspE* | -2.38 | -1.25 | 4.11E-06 | Transcription antiterminator |
| *eamB* | -2.38 | -1.25 | 6.23E-05 | Cysteine/O-acetylserine exporter |
| *pdeR* | -2.38 | -1.25 | 4.58E-06 | Cyclic di-GMP phosphodiesterase |
| *fic* | -2.37 | -1.24 | 5.28E-06 | Putative adenosine monophosphate transferase |
| *ybeD* | -2.37 | -1.24 | 1.67E-05 | DUF493 domain-containing protein |
| *aspA* | -2.36 | -1.24 | 4.77E-06 | Aspartase |

| | | | | |
|---|---|---|---|---|
| *clpB* | -2.36 | -1.24 | 5.12E-06 | Heat shock protein |
| *lpp* | -2.36 | -1.24 | 4.66E-06 | Murein lipoprotein |
| *rpmE* | -2.36 | -1.24 | 5.44E-06 | 50S ribosomal subunit protein L31 |
| *yjcH* | -2.36 | -1.24 | 6.44E-04 | DUF485 domain-containing inner membrane protein |
| *frdA* | -2.35 | -1.23 | 6.03E-06 | Fumarate reductase flavoprotein subunit |
| *ybdD* | -2.35 | -1.23 | 2.12E-05 | PF04328 family protein |
| *yedK* | -2.35 | -1.23 | 1.33E-05 | Genome maintenance protein |
| *yjaB* | -2.35 | -1.23 | 2.50E-05 | Peptidyl-lysine N-acetyltransferase |
| *loiP* | -2.34 | -1.23 | 8.84E-06 | Metalloprotease |
| *ysaB* | -2.34 | -1.23 | 2.21E-03 | Putative lipoprotein |
| *YjjY* | -2.33 | -1.22 | 8.41E-06 | Uncharacterised protein |
| *Z4874* | -2.33 | -1.22 | 1.01E-05 | Putative regulator |
| *cysQ* | -2.32 | -1.21 | 8.21E-06 | 3'(2'),5'-bisphosphate nucleotidase |
| *yjbE* | -2.32 | -1.21 | 9.06E-06 | Hypothetical protein |
| *Z5148* | -2.32 | -1.21 | 2.76E-04 | Hypothetical protein |
| *bioD* | -2.31 | -1.21 | 1.77E-05 | Dethiobiotin synthetase |
| *ucpA* | -2.31 | -1.21 | 9.94E-06 | Oxidoreductase |
| *ybeZ* | -2.31 | -1.21 | 9.35E-06 | PhoH-like protein |
| *yphA* | -2.31 | -1.21 | 5.16E-05 | Hypothetical protein |
| *ynjH* | -2.3 | -1.20 | 4.78E-04 | DUF1496 domain-containing protein |
| *Z_RS33335* | -2.3 | -1.20 | 9.56E-06 | Hypothetical protein |
| *ilvB* | -2.29 | -1.20 | 2.08E-05 | Acetohydroxy acid synthase I subunit |
| *mcbR* | -2.29 | -1.20 | 2.99E-05 | DNA-binding transcriptional dual regulator |
| *ssrS* | -2.29 | -1.20 | 1.14E-05 | 6S RNA |
| *agaD* | -2.28 | -1.19 | 0.01 | Galactosamine-specific PTS enzyme IID component |
| *cecR* | -2.28 | -1.19 | 5.23E-05 | DNA-binding transcriptional dual regulator |
| *dadA* | -2.28 | -1.19 | 1.33E-05 | D-amino acid dehydrogenase |
| *nagE* | -2.28 | -1.19 | 1.73E-05 | N-acetylglucosamine-specific PTS IIABC component |
| *yjbJ* | -2.28 | -1.19 | 1.23E-05 | Putative stress response protein |
| *cnoX* | -2.27 | -1.18 | 2.28E-05 | Chaperedoxin |
| *gutQ* | -2.27 | -1.18 | 4.67E-05 | D-arabinose 5-phosphate isomerase |
| *rseA* | -2.27 | -1.18 | 1.53E-05 | Anti-sigma-E factor |
| *ushA* | -2.27 | -1.18 | 1.78E-05 | 5'-nucleotidase / UDP-sugar hydrolase |
| *Z2302* | -2.27 | -1.18 | 1.54E-04 | Unknown protein encoded within prophage CP-933U |
| *ffs* | -2.26 | -1.18 | 2.50E-05 | 4.5S RNA |
| *nagB* | -2.26 | -1.18 | 2.47E-05 | Glucosamine-6-phosphate deaminase |
| *speA* | -2.26 | -1.18 | 1.84E-05 | Biosynthetic arginine decarboxylase |
| *Z2149* | -2.26 | -1.18 | 9.21E-04 | Hypothetical protein |
| *hslR* | -2.25 | -1.17 | 1.79E-04 | Heat shock protein Hsp15 |
| *ycgN* | -2.25 | -1.17 | 5.04E-05 | Putative metal-chelating domain-containing protein |
| *yncL* | -2.25 | -1.17 | 2.19E-05 | Stress response membrane protein |
| *hcaR* | -2.24 | -1.16 | 4.61E-05 | DNA-binding transcriptional dual regulator |
| *rrf* | -2.23 | -1.16 | 2.54E-05 | 5S ribosomal RNA |
| *ybaB* | -2.23 | -1.16 | 3.48E-05 | Putative nucleoid-associated protein |
| *ygeW* | -2.23 | -1.16 | 6.72E-03 | Putative carbamoyltransferase |
| *yihI* | -2.23 | -1.16 | 3.15E-05 | Der GTPase-activating protein |
| *yjfP* | -2.23 | -1.16 | 8.00E-05 | Carboxylesterase |
| *gpr* | -2.22 | -1.15 | 3.68E-05 | L-glyceraldehyde 3-phosphate reductase |
| *tas* | -2.22 | -1.15 | 4.91E-05 | NADP(H)-dependent aldo-keto reductase |
| *fdnI* | -2.21 | -1.14 | 2.09E-03 | Formate dehydrogenase N subunit γ |
| *frdB* | -2.21 | -1.14 | 4.16E-05 | Fumarate reductase iron-sulfur protein |

| | | | | |
|---|---|---|---|---|
| *yncJ* | -2.21 | -1.14 | 7.05E-04 | DUF2554 domain-containing protein |
| *Z0609* | -2.21 | -1.14 | 3.03E-05 | Hypothetical protein |
| *cybC* | -2.2 | -1.14 | 6.51E-05 | Cytochrome b (562) |
| *osmF* | -2.2 | -1.14 | 3.65E-04 | Glycine betaine ABC transporter periplasmic binding |
| *ybaA* | -2.2 | -1.14 | 9.97E-05 | DUF1428 domain-containing protein |
| *yhcO* | -2.2 | -1.14 | 5.16E-05 | Putative barnase inhibitor |
| *Z5890* | -2.2 | -1.14 | 4.64E-05 | Partial putative integrase |
| *dapE* | -2.19 | -1.13 | 4.77E-05 | Succinyl-diaminopimelate desuccinylase |
| *decR* | -2.19 | -1.13 | 1.33E-04 | DNA-binding transcriptional activator |
| *nnr* | -2.19 | -1.13 | 5.74E-05 | NAD(P)HX epimerase / NAD(P)HX dehydratase |
| *yafN* | -2.19 | -1.13 | 4.07E-04 | Antitoxin |
| *yicH* | -2.19 | -1.13 | 5.06E-05 | AsmA family protein |
| *Z0510* | -2.19 | -1.13 | 5.16E-05 | Hypothetical protein |
| *Z0608* | -2.19 | -1.13 | 4.22E-05 | Putative outer membrane export protein |
| *yceM* | -2.17 | -1.12 | 7.88E-05 | Putative oxidoreductase |
| *Z6057* | -2.17 | -1.12 | 1.29E-03 | tRNA-Arg |
| *degP* | -2.16 | -1.11 | 7.05E-05 | Periplasmic serine endoprotease |
| *dmlR* | -2.16 | -1.11 | 1.59E-04 | DNA-binding transcriptional regulator |
| *ecnB* | -2.16 | -1.11 | 6.05E-05 | Entericidin B |
| *hokE_1* | -2.16 | -1.11 | 3.66E-04 | Type I toxin-antitoxin system toxin |
| *csrA* | -2.15 | -1.10 | 1.98E-04 | Carbon storage regulato |
| *macB* | -2.15 | -1.10 | 8.32E-05 | ABC-type tripartite efflux pump ATP binding |
| *mtlA* | -2.15 | -1.10 | 6.59E-05 | Mannitol-specific PTS enzyme IICBA component |
| *pdeB* | -2.15 | -1.10 | 7.32E-05 | c-di-GMP phosphodiesterase |
| *rseD* | -2.15 | -1.10 | 1.31E-04 | RpoE leader peptide |
| *yajL* | -2.15 | -1.10 | 1.28E-04 | Protein/nucleic acid deglycase 3 |
| *yedI* | -2.15 | -1.10 | 1.79E-04 | DUF808 domain-containing inner membrane protein |
| *yneJ* | -2.15 | -1.10 | 4.50E-04 | Putative DNA-binding transcriptional regulator |
| *Z1874* | -2.15 | -1.10 | 0.03 | Putative antiterminator Q of prophage CP-933X |
| *mscK* | -2.14 | -1.10 | 9.69E-05 | Potassium dependent mechanosensitive channel |
| *nadR* | -2.14 | -1.10 | 8.49E-05 | DNA-binding transcriptional repressor |
| *ygiN* | -2.14 | -1.10 | 9.88E-05 | Hypothetical protein |
| *hokD_5* | -2.13 | -1.09 | 1.04E-04 | Type I toxin-antitoxin system toxin |
| *tnaA* | -2.13 | -1.09 | 6.33E-04 | Tryptophanase |
| *lysP* | -2.12 | -1.08 | 1.45E-04 | lysine:H+ symporter |
| *gatZ* | -2.11 | -1.08 | 0.03 | Putative tagatose-1,6-bisphosphate chaperone |
| *sfsA* | -2.11 | -1.08 | 1.01E-04 | Sugar fermentation stimulation protein A |
| *ygiB* | -2.11 | -1.08 | 1.22E-04 | DUF1190 domain-containing protein |
| *blc* | -2.1 | -1.07 | 1.40E-04 | Outer membrane lipoprotein |
| *dksA* | -2.1 | -1.07 | 1.23E-04 | RNA polymerase-binding transcription factor |
| *hslV* | -2.1 | -1.07 | 1.77E-04 | Peptidase component of the HslVU protease |
| *yahO* | -2.1 | -1.07 | 1.16E-04 | DUF1471 domain-containing protein |
| *adiY* | -2.09 | -1.06 | 1.16E-03 | DNA-binding transcriptional activator |
| *asr* | -2.09 | -1.06 | 3.65E-03 | Periplasmic chaperon |
| *slyB* | -2.09 | -1.06 | 1.38E-04 | Outer membrane lipoprotein |
| *uxuR* | -2.09 | -1.06 | 1.79E-04 | DNA-binding transcriptional repressor |
| *agaE* | -2.08 | -1.06 | 0.01 | Putative phosphotransferase system enzyme subunit |
| *cysK* | -2.08 | -1.06 | 1.41E-04 | Cysteine synthase A |
| *ypeB* | -2.08 | -1.06 | 6.90E-04 | PF12843 family protein |
| *Z_RS09840* | -2.08 | -1.06 | 3.24E-04 | Hok/Gef family protein |
| *marC* | -2.07 | -1.05 | 4.40E-04 | Inner membrane protein |

| | | | | |
|---|---|---|---|---|
| *narK* | -2.07 | -1.05 | 4.09E-03 | Nitrite extrusion protein |
| *sufA* | -2.07 | -1.05 | 2.23E-04 | Fe-S cluster assembly scaffold |
| *yehP* | -2.07 | -1.05 | 0.03 | VWA domain-containing protein |
| *cdd* | -2.06 | -1.04 | 2.05E-04 | Cytidine/deoxycytidine deaminase |
| *eptA* | -2.06 | -1.04 | 1.85E-03 | Phosphoethanolamine transferase |
| *sixA* | -2.06 | -1.04 | 1.75E-04 | Phosphohistidine phosphatase |
| *yeaQ* | -2.06 | -1.04 | 2.09E-04 | PF04226 family protein |
| *potE* | -2.05 | -1.04 | 1.18E-03 | Putrescine transport protein |
| *rsd* | -2.05 | -1.04 | 2.63E-04 | Regulator of sigma D |
| *ybbW* | -2.05 | -1.04 | 2.17E-03 | Putative allantoin permease |
| *Z4776* | -2.05 | -1.04 | 3.99E-04 | Hypothetical protein |
| *ldtA* | -2.04 | -1.03 | 2.38E-04 | L,D-transpeptidase |
| *mlaC* | -2.04 | -1.03 | 4.00E-04 | Intermembrane phospholipid transport system |
| *uspF* | -2.04 | -1.03 | 2.14E-04 | Universal stress protein F |
| *Z4882* | -2.04 | -1.03 | 3.00E-04 | Type II toxin-antitoxin system HicB family antitoxin |
| *Z6073* | -2.04 | -1.03 | 2.23E-04 | Putative repressor cryptic prophage CP-933P |
| *arnT* | -2.03 | -1.02 | 6.35E-04 | Lipid IVA 4-amino-4-deoxy-L-arabinosyltransferase |
| *frwB* | -2.03 | -1.02 | 2.75E-03 | PTS system fructose-like IIB component 1 |
| *ldcC* | -2.03 | -1.02 | 2.97E-04 | Lysine decarboxylase 2 |
| *nifJ* | -2.03 | -1.02 | 2.57E-04 | Flavodoxin oxidoreductase |
| *ybiB* | -2.03 | -1.02 | 3.01E-04 | Non-specific DNA-binding protein |
| *Z1383* | -2.03 | -1.02 | 2.84E-03 | Unknown protein, cryptic prophage CP-933M |
| *Z3390* | -2.03 | -1.02 | 2.95E-04 | Putative hydroxylase |
| *yfcG* | -2.02 | -1.01 | 5.91E-04 | Disulphide bond oxidoreductase |
| *Z_RS29520* | -2.02 | -1.01 | 2.66E-03 | KGG domain-containing protein |
| *ghoS_2* | -2.01 | -1.01 | 2.90E-03 | Antitoxin of the GhoTS toxin-antitoxin system |
| *grpE* | -2.01 | -1.01 | 3.74E-04 | Nucleotide exchange factor |
| *rpoE* | -2.01 | -1.01 | 3.42E-04 | RNA polymerase, sigma-E factor |
| *rtcB* | -2.01 | -1.01 | 1.45E-03 | Orf, hypothetical protein |
| *yghA* | -2.01 | -1.01 | 6.50E-04 | NADP+-dependent aldehyde reductase |
| *ppnN* | -2 | -1.00 | 3.56E-04 | Nucleotide 5'-monophosphate nucleosidase |
| *yeeE* | -2 | -1.00 | 6.24E-04 | Thiosulphate transporter |
| *Z4324* | -2 | -1.00 | 3.76E-03 | Putative transposase |
| *cusR* | -1.99 | -0.99 | 2.83E-03 | DNA-binding transcriptional activator |
| *hokD_1* | -1.99 | -0.99 | 5.33E-04 | Type I toxin-antitoxin system toxin |
| *rnd* | -1.99 | -0.99 | 7.96E-04 | RNase D |
| *Z0347* | -1.99 | -0.99 | 8.10E-03 | Hypothetical protein |
| *Z5619* | -1.99 | -0.99 | 3.56E-03 | Putative transcriptional regulator of sorbose uptake |
| *fucA* | -1.98 | -0.99 | 2.54E-03 | L-fuculose-1-phosphate aldolase |
| *dkgB* | -1.97 | -0.98 | 1.77E-03 | Methylglyoxal reductase |
| *gloB* | -1.97 | -0.98 | 7.28E-04 | Hydroxyacylglutathione hydrolase |
| *ycfH* | -1.97 | -0.98 | 5.47E-04 | Putative metal-dependent hydrolase |
| *yciW* | -1.97 | -0.98 | 0.01 | Putative peroxidase |
| *ydiL* | -1.97 | -0.98 | 0.02 | DUF1870 domain-containing protein |
| *yfgG* | -1.97 | -0.98 | 4.96E-04 | Nickel/cobalt stress response protein |
| *yhjD* | -1.97 | -0.98 | 5.15E-04 | Putative transporter |
| *ompR* | -1.96 | -0.97 | 6.62E-04 | DNA-binding dual transcriptional regulator |
| *yajI* | -1.96 | -0.97 | 6.20E-03 | Hypothetical protein |
| *ycfP* | -1.96 | -0.97 | 6.57E-04 | PF05728 family protein |
| *yeaW* | -1.96 | -0.97 | 0.01 | Carnitine monooxygenase subunit |
| *arcA* | -1.95 | -0.96 | 6.55E-04 | DNA-binding transcriptional dual regulator |

| | | | | |
|---|---|---|---|---|
| *ligA* | -1.95 | -0.96 | 1.35E-03 | DNA ligase |
| *trkA* | -1.95 | -0.96 | 1.15E-03 | NAD-binding component of Trk potassium transporters |
| *uspA* | -1.95 | -0.96 | 6.33E-04 | Universal stress protein A |
| *ytfE* | -1.95 | -0.96 | 6.22E-04 | Iron-sulphur cluster repair protein |
| *gcvT* | -1.94 | -0.96 | 5.09E-03 | Aminomethyltransferase |
| *hlyD* | -1.94 | -0.96 | 8.86E-04 | Secretion protein |
| *mdaB* | -1.94 | -0.96 | 3.58E-03 | NADPH:quinone oxidoreductase |
| *yibH* | -1.94 | -0.96 | 1.45E-03 | Inner membrane protein |
| *yjdC* | -1.94 | -0.96 | 9.26E-04 | Putative DNA-binding transcriptional regulator |
| *crr* | -1.93 | -0.95 | 8.07E-04 | PTS system, glucose-specific IIA component |
| *ihfB* | -1.93 | -0.95 | 8.99E-04 | Integration host factor subunit β |
| *melA* | -1.93 | -0.95 | 1.58E-03 | Alpha-galactosidase |
| *ydeN* | -1.93 | -0.95 | 2.07E-03 | Putative sulfatase |
| *yncG* | -1.93 | -0.95 | 0.03 | Putative glutathione S-transferase |
| *nagA* | -1.92 | -0.94 | 1.18E-03 | N-acetylglucosamine-6-phosphate deacetylase |
| *pheT* | -1.92 | -0.94 | 9.08E-04 | Phenylalanine tRNA synthetase, beta-subunit |
| *rssB* | -1.92 | -0.94 | 1.06E-03 | Two-component system response regulator |
| *ybaE* | -1.92 | -0.94 | 1.16E-03 | Putative nucleoid-associated protein |
| *yjbB* | -1.92 | -0.94 | 1.91E-03 | Putative inorganic phosphate export protein |
| *acpP* | -1.91 | -0.93 | 9.91E-04 | Acyl carrier protein |
| *emtA* | -1.91 | -0.93 | 2.09E-03 | Murein transglycosylase E |
| *higA* | -1.91 | -0.93 | 3.65E-03 | Antitoxin/DNA-binding transcriptional repressor |
| *nudC* | -1.91 | -0.93 | 2.46E-03 | RNA decapping hydrolase |
| *ypfJ* | -1.91 | -0.93 | 1.15E-03 | Uncharacterised protein |
| *zitB* | -1.91 | -0.93 | 1.77E-03 | Zn2+/Cd2+/Ni2+/Cu2+ exporter |
| *aldA* | -1.9 | -0.93 | 1.15E-03 | Aldehyde dehydrogenase |
| *sstT* | -1.9 | -0.93 | 1.09E-03 | Serine/threonine transporter |
| *ybjQ* | -1.9 | -0.93 | 2.39E-03 | Putative heavy metal binding protein |
| *yegP* | -1.9 | -0.93 | 1.15E-03 | Hypothetical protein |
| *yifK* | -1.9 | -0.93 | 6.35E-03 | Putative transporter |
| *zupT* | -1.9 | -0.93 | 1.96E-03 | Divalent metal ion transporter |
| *cysA* | -1.89 | -0.92 | 3.34E-03 | Sulphate/thiosulphate ABC transporter ATP subunit |
| *rpoH* | -1.89 | -0.92 | 1.40E-03 | RNA polymerase, sigma(32) factor |
| *ynfL* | -1.89 | -0.92 | 0.01 | Putative DNA-binding transcriptional regulator |
| *gmhB* | -1.88 | -0.91 | 1.64E-03 | D-glycero-β-D-manno-heptose-1,7-bisphosphate |
| *ibpA* | -1.88 | -0.91 | 2.16E-03 | Small heat shock protein |
| *msyB_1* | -1.88 | -0.91 | 1.53E-03 | Acidic protein |
| *queE* | -1.88 | -0.91 | 1.53E-03 | Putative 7-carboxy-7-deazaguanine synthase |
| *sbp* | -1.88 | -0.91 | 0.01 | Periplasmic sulfate-binding protein |
| *yggE* | -1.88 | -0.91 | 1.79E-03 | DUF541 domain-containing protein |
| *yidB* | -1.88 | -0.91 | 1.45E-03 | DUF937 domain-containing protein |
| *Z0972* | -1.88 | -0.91 | 8.68E-03 | Putative tail component of prophage CP-933K |
| *dnaJ* | -1.87 | -0.90 | 1.74E-03 | Chaperone protein |
| *katG* | -1.87 | -0.90 | 1.98E-03 | Catalase |
| *pflA* | -1.87 | -0.90 | 1.98E-03 | Pyruvate formate lyase activating enzyme 1 |
| *ybbJ* | -1.87 | -0.90 | 1.77E-03 | NfeD-like family protein |
| *yfbV* | -1.87 | -0.90 | 1.64E-03 | PF04217 family membrane protein |
| *Z_RS29535* | -1.87 | -0.90 | 0.02 | ACP S-malonyltransferase |
| *kbp* | -1.86 | -0.90 | 1.94E-03 | K+ binding protein |
| *mlc* | -1.86 | -0.90 | 3.07E-03 | Putative NAGC-like transcriptional regulator |
| *msyB_2* | -1.86 | -0.90 | 5.74E-03 | Acidic protein |

| | | | | |
|---|---|---|---|---|
| *sdaC* | -1.86 | -0.90 | 3.50E-03 | Probable serine transporter |
| *ybjD* | -1.86 | -0.90 | 2.09E-03 | DUF2813 domain-containing protein |
| *yiiM* | -1.86 | -0.90 | 1.91E-03 | 6-hydroxyaminopurine reductase |
| *hslO* | -1.85 | -0.89 | 3.16E-03 | Molecular chaperone Hsp33 |
| *nrdG* | -1.85 | -0.89 | 4.26E-03 | Anaerobic ribonucleotide reductase activating protein |
| *ygaP* | -1.85 | -0.89 | 5.35E-03 | Thiosulphate sulphurtransferase |
| *yhbO* | -1.85 | -0.89 | 4.80E-03 | Protein/nucleic acid deglycase 2 |
| *yijD* | -1.85 | -0.89 | 4.64E-03 | DUF1422 domain-containing inner membrane protein |
| *araC* | -1.84 | -0.88 | 2.42E-03 | Transcriptional regulator |
| *ldtC* | -1.84 | -0.88 | 5.28E-03 | L,D-transpeptidase |
| *napC* | -1.84 | -0.88 | 0.03 | Cytochrome c-type protein |
| *yihV* | -1.84 | -0.88 | 0.01 | 6-deoxy-6-sulfofructose kinase |
| *yohF* | -1.84 | -0.88 | 4.71E-03 | Putative oxidoreductase |
| *Z3118* | -1.84 | -0.88 | 0.01 | Unknown protein encoded within prophage CP-933U |
| *cmk* | -1.83 | -0.87 | 3.70E-03 | Cytidylate kinase |
| *cof* | -1.83 | -0.87 | 7.58E-03 | HMP-PP phosphatase |
| *cycA* | -1.83 | -0.87 | 2.51E-03 | D-serine/alanine/glycine/:H+symporter |
| *dgcM* | -1.83 | -0.87 | 4.64E-03 | Diguanylate cyclase |
| *mlaA* | -1.83 | -0.87 | 3.80E-03 | Intermembrane phospholipid transport system |
| *mlaF* | -1.83 | -0.87 | 3.57E-03 | Intermembrane phospholipid transport system |
| *mtlR* | -1.83 | -0.87 | 2.79E-03 | Transcriptional repressor |
| *prlF* | -1.83 | -0.87 | 4.48E-03 | Antitoxin |
| *ybhD* | -1.83 | -0.87 | 0.03 | Putative DNA-binding transcriptional regulator |
| *yiiR* | -1.83 | -0.87 | 0.02 | DUF805 domain-containing protein |
| *Z3123* | -1.83 | -0.87 | 4.09E-03 | Unknown protein encoded within prophage CP-933U |
| *chaB* | -1.82 | -0.86 | 5.25E-03 | Putative cation transport regulator |
| *dtpB* | -1.82 | -0.86 | 3.00E-03 | Dipeptide/tripeptide:H+ symporter |
| *inaA* | -1.82 | -0.86 | 7.95E-03 | Putative lipopolysaccharide kinase |
| *otsB* | -1.82 | -0.86 | 3.77E-03 | Trehalose-6-phosphate phophatase |
| *yihO* | -1.82 | -0.86 | 0.02 | Putative sulfoquinovose transporter |
| *yjgM* | -1.82 | -0.86 | 6.09E-03 | GNAT family N-acetyltransferase |
| *yphF* | -1.82 | -0.86 | 0.02 | Putative ABC transporter periplasmic binding protein |
| *fxsA* | -1.81 | -0.86 | 4.34E-03 | Hypothetical protein |
| *hns* | -1.81 | -0.86 | 2.80E-03 | DNA-binding transcriptional dual regulator |
| *rpnB* | -1.81 | -0.86 | 4.52E-03 | Recombination-promoting nuclease |
| *rraB* | -1.81 | -0.86 | 3.65E-03 | Ribonuclease E inhibitor protein B |
| *sufB* | -1.81 | -0.86 | 3.27E-03 | Fe-S cluster assembly protein |
| *tus* | -1.81 | -0.86 | 4.60E-03 | DNA replication terminus site-binding protein |
| *yaiY* | -1.81 | -0.86 | 8.78E-03 | DUF2755 domain-containing inner membrane protein |
| *ydhQ* | -1.81 | -0.86 | 3.93E-03 | Putative adhesin-related protein |
| *ysgA* | -1.81 | -0.86 | 3.40E-03 | Putative dienelactone hydrolase |
| *frr* | -1.8 | -0.85 | 3.66E-03 | Ribosome recycling factor |
| *tisB* | -1.8 | -0.85 | 0.02 | Type I toxin-antitoxin system toxin |
| *Z4787* | -1.8 | -0.85 | 0.02 | Hypothetical protein |
| *astC* | -1.79 | -0.84 | 3.68E-03 | Succinylornithine transaminase |
| *hcr* | -1.79 | -0.84 | 0.03 | NADH oxidoreductase |
| *pnuC* | -1.79 | -0.84 | 7.55E-03 | Nicotinamide riboside transporter |
| *rimI* | -1.79 | -0.84 | 0.02 | N-acetyltransferase |
| *sapA* | -1.79 | -0.84 | 4.35E-03 | Putative periplasmic binding protein |
| *tnaB* | -1.79 | -0.84 | 0.03 | Low affinity tryptophan permease |
| *yaeH* | -1.79 | -0.84 | 4.54E-03 | DUF3461 domain-containing protein |

| | | | | |
|---|---|---|---|---|
| *yeaV* | -1.79 | -0.84 | 0.02 | Putative transporter |
| *adeD* | -1.78 | -0.83 | 7.19E-03 | Adenine deaminase |
| *bamB* | -1.78 | -0.83 | 4.60E-03 | Outer membrane protein assembly factor |
| *hokE_2* | -1.78 | -0.83 | 0.01 | Type I toxin-antitoxin system toxin |
| *holE* | -1.78 | -0.83 | 0.03 | DNA polymerase III subunit θ |
| *ppsA* | -1.78 | -0.83 | 4.09E-03 | Phosphoenolpyruvate synthase |
| *puuR* | -1.78 | -0.83 | 0.02 | DNA-binding transcriptional repressor |
| *yafV* | -1.78 | -0.83 | 8.83E-03 | Antitoxin |
| *yecN* | -1.78 | -0.83 | 7.22E-03 | MAPEG family inner membrane protein |
| *Z0056* | -1.78 | -0.83 | 7.45E-03 | Putative antitoxin of gyrase inhibiting toxin-antitoxin |
| *zapA* | -1.78 | -0.83 | 4.59E-03 | Cell division protein |
| *iaaA* | -1.77 | -0.82 | 5.84E-03 | β-aspartyl-peptidase |
| *rpmI* | -1.77 | -0.82 | 4.93E-03 | 50S ribosomal subunit protein A |
| *ydgD* | -1.77 | -0.82 | 4.50E-03 | Putative serine protease |
| *yegS* | -1.77 | -0.82 | 5.91E-03 | Lipid kinase |
| *nrdH* | -1.76 | -0.82 | 5.71E-03 | Glutaredoxin-like protein |
| *queG* | -1.76 | -0.82 | 8.18E-03 | Epoxyqueuosine reductase |
| *Z_RS08560* | -1.76 | -0.82 | 0.01 | AlpA family transcriptional regulator |
| *Z1769* | -1.76 | -0.82 | 6.45E-03 | Unknown protein prophage CP-933N |
| *Z6074* | -1.76 | -0.82 | 5.77E-03 | Unknown protein cryptic prophage CP-933P |
| *fmt* | -1.75 | -0.81 | 5.89E-03 | N-formyltransferase |
| *fnr* | -1.75 | -0.81 | 6.09E-03 | DNA-binding transcriptional dual regulator |
| *fucO* | -1.75 | -0.81 | 6.01E-03 | L-1,2-propanediol oxidoreductase |
| *opgD* | -1.75 | -0.81 | 8.05E-03 | Glucans biosynthesis protein D |
| *Z1766* | -1.75 | -0.81 | 7.43E-03 | Unknown protein encoded by prophage CP-933N |
| *ackA* | -1.74 | -0.80 | 6.58E-03 | Acetate kinase |
| *ftsX* | -1.74 | -0.80 | 7.74E-03 | Cell division membrane protein |
| *uxaB* | -1.74 | -0.80 | 8.40E-03 | Tagaturonate reductase |
| *chbC* | -1.73 | -0.79 | 0.04 | N,N'-diacetylchitobiose PTS enzyme IIC component |
| *espM1* | -1.73 | -0.79 | 7.43E-03 | T3SS effector |
| *sapB* | -1.73 | -0.79 | 9.79E-03 | Putrescine ABC exporter membrane subunit |
| *yahN* | -1.73 | -0.79 | 9.70E-03 | 2-oxoglutaramate amidase |
| *ybhR* | -1.73 | -0.79 | 0.01 | ABC exporter membrane subunit |
| *ygdR* | -1.73 | -0.79 | 8.83E-03 | DUF903 domain-containing lipoprotein |
| *yqjA* | -1.73 | -0.79 | 8.12E-03 | DedA family protein |
| *Z1345* | -1.73 | -0.79 | 0.01 | Qin prophage; putative antitermination protein Q |
| *Z1768* | -1.73 | -0.79 | 7.02E-03 | Unknown protein encoded by prophage CP-933N |
| *Z4883* | -1.73 | -0.79 | 0.01 | Type II toxin-antitoxin system HicA family toxin |
| *nanA* | -1.72 | -0.78 | 0.02 | N-acetylneuraminate lyase |
| *yccT* | -1.72 | -0.78 | 8.29E-03 | DUF2057 domain-containing protein |
| *yebG* | -1.72 | -0.78 | 7.86E-03 | DNA damage-inducible protein |
| *yqgA* | -1.72 | -0.78 | 0.02 | DUF554 domain-containing protein |
| *Z_RS08615* | -1.72 | -0.78 | 0.02 | PerC family transcriptional regulator |
| *Z1781* | -1.72 | -0.78 | 8.30E-03 | Unknown protein encoded by prophage CP-933N |
| *Z4070* | -1.72 | -0.78 | 0.01 | CRISPR-associated helicase/endonuclease Cas3 |
| *cysC* | -1.71 | -0.77 | 0.04 | Adenylyl-sulphate kinase |
| *idi* | -1.71 | -0.77 | 0.01 | Isopentenyl-diphosphate Δ-isomerase |
| *ygjG* | -1.71 | -0.77 | 8.78E-03 | Putrescine aminotransferase |
| *Z0967* | -1.71 | -0.77 | 0.03 | Putative protease encoded in prophage CP-933K |
| *Z1765* | -1.71 | -0.77 | 0.01 | Putative excisionase for prophage CP-933N |
| *allE* | -1.7 | -0.77 | 0.03 | (S)-ureidoglycine aminohydrolase |

| | | | | |
|---|---|---|---|---|
| *anmK* | -1.7 | -0.77 | 0.02 | Anhydro-N-acetylmuramic acid kinase |
| *rna* | -1.7 | -0.77 | 0.01 | RNase I |
| *tdcG* | -1.7 | -0.77 | 0.05 | L-serine ammonia-lyase |
| *yebV* | -1.7 | -0.77 | 9.61E-03 | DUF1480 domain-containing protein |
| *yqhC* | -1.7 | -0.77 | 0.02 | Putative ARAC-type regulatory |
| *ytfB* | -1.7 | -0.77 | 0.01 | Cell division protein |
| *ascB* | -1.69 | -0.76 | 0.02 | 6-phospho-β-glucosidase |
| *betI* | -1.69 | -0.76 | 0.02 | DNA-binding transcriptional repressor |
| *garD* | -1.69 | -0.76 | 0.02 | Galactarate dehydratase |
| *garK* | -1.69 | -0.76 | 0.02 | Glycerate 2-kinase 1 |
| *malM* | -1.69 | -0.76 | 0.01 | Maltose regulon periplasmic protein |
| *sra* | -1.69 | -0.76 | 0.01 | Ribosome-associated protein |
| *ydcF* | -1.69 | -0.76 | 0.02 | DUF218 domain-containing protein |
| *ydhF* | -1.69 | -0.76 | 0.01 | Hypothetical protein |
| *yeeZ* | -1.69 | -0.76 | 0.01 | Putative epimerase |
| *yeiE* | -1.69 | -0.76 | 0.01 | DNA-binding transcriptional dual regulator |
| *yfdE* | -1.69 | -0.76 | 0.03 | Hypothetical protein |
| *yhbW* | -1.69 | -0.76 | 0.02 | Putative luciferase-like monooxygenase |
| *yhbX* | -1.69 | -0.76 | 0.02 | Putative hydrolase |
| *Z0266* | -1.69 | -0.76 | 0.02 | Hypothetical protein |
| *fur* | -1.68 | -0.75 | 0.01 | Negative regulator |
| *glnB* | -1.68 | -0.75 | 0.03 | Nitrogen regulatory protein PII |
| *hcaT* | -1.68 | -0.75 | 0.01 | Putative 3-phenylpropionate transporter |
| *hokD_2* | -1.68 | -0.75 | 0.01 | Type I toxin-antitoxin system toxin |
| *mocA* | -1.68 | -0.75 | 0.01 | Molybdenum cofactor cytidylyltransferase |
| *oppF* | -1.68 | -0.75 | 0.03 | Murein tripeptide ABC transporter |
| *pfkB* | -1.68 | -0.75 | 0.01 | 6-phosphofructokinase II |
| *yhaV* | -1.68 | -0.75 | 0.01 | Ribosome-dependent mRNA interferase toxin |
| *Z_RS32620* | -1.68 | -0.75 | 0.02 | NadS family protein |
| *Z1772* | -1.68 | -0.75 | 0.01 | Unknown protein encoded by prophage CP-933N |
| *espZ* | -1.67 | -0.74 | 0.02 | T3SS effector |
| *hslU* | -1.67 | -0.74 | 0.01 | ATPase component of the HslVU protease |
| *pmbA* | -1.67 | -0.74 | 0.02 | Metalloprotease subunit |
| *recN* | -1.67 | -0.74 | 0.01 | DNA repair protein |
| *speB* | -1.67 | -0.74 | 0.02 | Agmatinase |
| *Z1776* | -1.67 | -0.74 | 0.01 | Unknown protein encoded by prophage CP-933N |
| *Z5151* | -1.67 | -0.74 | 0.02 | Hypothetical protein |
| *gntP* | -1.66 | -0.73 | 0.03 | Fructuronate transporter |
| *kdgT* | -1.66 | -0.73 | 0.04 | 2-dehydro-3-deoxy-D-gluconate:H+symporte |
| *metL* | -1.66 | -0.73 | 0.02 | Aspartokinase II and homoserine dehydrogenase II |
| *yaaU* | -1.66 | -0.73 | 0.02 | Putative transporter |
| *yccM* | -1.66 | -0.73 | 0.04 | Putative electron transport protein |
| *yjeH* | -1.66 | -0.73 | 0.02 | L-methionine/branched chain amino acid exporter |
| *yrdA* | -1.66 | -0.73 | 0.02 | Putative transferase |
| *zntR* | -1.66 | -0.73 | 0.02 | DNA-binding transcriptional activator |
| *ataT* | -1.65 | -0.72 | 0.02 | Hypothetical protein |
| *brnQ* | -1.65 | -0.72 | 0.02 | Branched chain amino acid transporter |
| *nanM* | -1.65 | -0.72 | 0.02 | N-acetylneuraminate mutarotase |
| *rimK* | -1.65 | -0.72 | 0.03 | Ribosomal protein S6 modification protein |
| *ybiU* | -1.65 | -0.72 | 0.03 | DUF1479 domain-containing protein |
| *ydcU* | -1.65 | -0.72 | 0.05 | Putative ABC transporter membrane subunit |

| | | | | |
|---|---|---|---|---|
| *yjdN* | -1.65 | -0.72 | 0.02 | PF06983 family protein |
| *yqjC* | -1.65 | -0.72 | 0.02 | DUF1090 domain-containing protein |
| *ytfF* | -1.65 | -0.72 | 0.03 | Inner membrane protein |
| *Z0956* | -1.65 | -0.72 | 0.02 | Putative antiterminator Q, prophage CP-933K |
| *Z1773* | -1.65 | -0.72 | 0.02 | Unknown protein encoded by prophage CP-933N |
| *flgK* | -1.64 | -0.71 | 0.04 | Flagellar hook-filament junction protein 1 |
| *hokD_4* | -1.64 | -0.71 | 0.02 | Type I toxin-antitoxin system toxin |
| *pepT* | -1.64 | -0.71 | 0.02 | Putative peptidase T |
| *ydcH* | -1.64 | -0.71 | 0.02 | Hypothetical protein |
| *yeaO* | -1.64 | -0.71 | 0.03 | DUF488 domain-containing protein |
| *yeeA* | -1.64 | -0.71 | 0.02 | Putative transporter |
| *Z1778* | -1.64 | -0.71 | 0.02 | Unknown protein encoded by prophage CP-933N |
| *Z3230* | -1.64 | -0.71 | 0.03 | Hypothetical protein |
| *dcuA* | -1.63 | -0.70 | 0.02 | C4-dicarboxylate transporter |
| *ldcA* | -1.63 | -0.70 | 0.03 | Murein tetrapeptide carboxypeptidase |
| *usg* | -1.63 | -0.70 | 0.02 | Putative semialdehyde dehydrogenase |
| *ycgM* | -1.63 | -0.70 | 0.02 | Putative isomerase/hydrolas |
| *ycjX* | -1.63 | -0.70 | 0.03 | DUF463 domain-containing protein |
| *yfhM* | -1.63 | -0.70 | 0.02 | α2-macroglobulin |
| *Z_RS12145* | -1.63 | -0.70 | 0.02 | Type II toxin-antitoxin system RelE/ParE family |
| *Z1771* | -1.63 | -0.70 | 0.02 | Rac prophage; putative transcriptional regulator |
| *Z1774* | -1.63 | -0.70 | 0.02 | Unknown protein encoded by prophage CP-933N |
| *Z1775* | -1.63 | -0.70 | 0.02 | Unknown protein encoded by prophage CP-933N |
| *clpP* | -1.62 | -0.70 | 0.02 | ATP-dependent Clp protease proteolytic subunit |
| *curA* | -1.62 | -0.70 | 0.03 | NADPH-dependent curcumin reductase |
| *rplT* | -1.62 | -0.70 | 0.02 | 50S ribosomal subunit protein L20 |
| *rraA* | -1.62 | -0.70 | 0.03 | Ribonuclease E inhibitor protein A |
| *sapC* | -1.62 | -0.70 | 0.03 | Peptide ABC transporter permease |
| *yceK* | -1.62 | -0.70 | 0.02 | DUF1375 domain-containing lipoprotein |
| *miaA* | -1.61 | -0.69 | 0.02 | Delta(2)-isopentenylpyrophosphate tRNA-transferase |
| *yaeP* | -1.61 | -0.69 | 0.03 | Hypothetical protein |
| *ycbJ* | -1.61 | -0.69 | 0.04 | Putative phosphotransferase |
| *ychN* | -1.61 | -0.69 | 0.03 | DsrE/F sulfur relay family protein |
| *yjjI* | -1.61 | -0.69 | 0.03 | DUF3029 domain-containing protein |
| *Z_RS08520* | -1.61 | -0.69 | 0.04 | T3SS effector |
| *Z_RS31570* | -1.61 | -0.69 | 0.04 | Hypothetical protein |
| *Z1764* | -1.61 | -0.69 | 0.02 | Partial integrase for prophage CP-933N |
| *dprA* | -1.6 | -0.68 | 0.03 | NAD(P)H-dependent nitroreductase |
| *elaB* | -1.6 | -0.68 | 0.03 | Tail-anchored inner membrane protein |
| *msrA* | -1.6 | -0.68 | 0.03 | Peptide methionine sulfoxide reductase |
| *sapD* | -1.6 | -0.68 | 0.03 | Putative ATP-binding protein of peptide transport |
| *treF* | -1.6 | -0.68 | 0.03 | Cytoplasmic trehalase |
| *astA* | -1.59 | -0.67 | 0.03 | Arginine N-succinyltransferase |
| *clpA* | -1.59 | -0.67 | 0.03 | Serine-protease ATP-binding component |
| *cyuA* | -1.59 | -0.67 | 0.03 | Putative L-cysteine desulfidase |
| *dnaG* | -1.59 | -0.67 | 0.03 | DNA primase |
| *espN* | -1.59 | -0.67 | 0.03 | T3SS effector |
| *hfq* | -1.59 | -0.67 | 0.03 | RNA binding protein |
| *kdgA* | -1.59 | -0.67 | 0.03 | KHG/KDPG aldolase |
| *metJ* | -1.59 | -0.67 | 0.05 | DNA-binding transcriptional repressor |
| *uxuA* | -1.59 | -0.67 | 0.03 | D-mannonate dehydratase |

| | | | | |
|---|---|---|---|---|
| *yaeR* | -1.59 | -0.67 | 0.04 | VOC domain-containing protein |
| *dedD* | -1.58 | -0.66 | 0.03 | Cell division membrane protein |
| *lgoR* | -1.58 | -0.66 | 0.03 | Hypothetical protein |
| *tal* | -1.58 | -0.66 | 0.03 | Transaldolase |
| *yicC* | -1.58 | -0.66 | 0.05 | Putative RNAse adapter protein |
| *bcr* | -1.57 | -0.65 | 0.04 | Multidrug efflux pump |
| *lldP* | -1.57 | -0.65 | 0.04 | Lactate/glycolate:H+ symporter |
| *sufC* | -1.57 | -0.65 | 0.04 | Fe-S cluster assembly ATPase |
| *ydcY* | -1.57 | -0.65 | 0.04 | DUF2526 domain-containing protein |
| *yohD* | -1.57 | -0.65 | 0.05 | DedA family protein |
| *Z1777* | -1.57 | -0.65 | 0.03 | Unknown protein encoded by prophage CP-933N |
| *amiD* | -1.56 | -0.64 | 0.04 | N-acetylmuramoyl-L-alanine amidase D |
| *espK* | -1.56 | -0.64 | 0.04 | T3SS effector |
| *gltS* | -1.56 | -0.64 | 0.04 | Glutamate:sodium symporter |
| *lpxC* | -1.56 | -0.64 | 0.04 | UDP-3-O-acyl N-acetylglucosamine deacetylase |
| *yidE* | -1.56 | -0.64 | 0.04 | Putative transport protein |
| *ybjP* | -1.55 | -0.63 | 0.04 | DUF3828 domain-containing lipoprotein |
| *ygeV* | -1.55 | -0.63 | 0.04 | Putative σ54-dependent transcriptional regulator |
| *ymdB* | -1.55 | -0.63 | 0.05 | 2'-O-acetyl-ADP-ribose deacetylase |
| *ada* | -1.54 | -0.62 | 0.05 | DNA-binding transcriptional dual regulator |
| *rlmE* | -1.54 | -0.62 | 0.04 | 23S rRNA 2'-O-ribose U2552 methyltransferase |
| *ybeY* | -1.54 | -0.62 | 0.05 | Endoribonuclease |
| *yihY* | -1.54 | -0.62 | 0.05 | PF03631 family membrane protein |

**Supplementary Table 2** – Bacterial strains used in this study

| Strain | Description | Reference |
|---|---|---|
| TUV93-0 | EHEC O157:H7 str. EDL933 $Stx^-$ | Campellone *et al*. 2002 |
| Δ*araC* | TUV93-0 *araC* deletion mutant; $Kan^R$ | This study |
| Δ*araBAD* | TUV93-0 *araBAD* deletion mutant; $Kan^R$ | This study |
| Δ*araFGH* | TUV93-0 *araFGH* deletion mutant; $Kan^R$ | This study |
| Δ*araE* | TUV93-0 *araE* deletion mutant; $Cm^R$ | This study |
| Δ*rbsDACBKR* | TUV93-0 *rbsDACBKR* deletion mutant; $Kan^R$ | This study |
| ZAP193 | EHEC O157:H7 str. ZAP193 (NCTC 12900) $Stx^-$ | Roe *et al.* 2004 |
| ZAP193 Δ*Z0415-9* | ZAP193 *Z0415-9* deletion mutant; $Kan^R$ $Strep^R$ | This study |
| ICC169 | *C. rodentium* O152 serotype; $Nal^R$ | Petty *et al.* 2010 |
| ICC169 Δ*araBAD* | ICC169 *araBAD* deletion mutant; $Kan^R$ | This study |
| ICC1370 | *C. rodentium* constitutive-$P_{ler}$ luminescence; $Nal^R$ $Kan^R$ | Mullineaux-Sanders *et al.* 2017 |
| ICC1370 Δ*araBAD* | *ICC1370 ΔaraBAD* mutant; $Nal^R$ $Kan^R$ $Cm^R$ | This study |

**Supplementary Table 2** – Plasmids used in this study

| **Plasmid** | **Description** | **Reference** |
|---|---|---|
| pMK1*lux* | pBR322 ori with the *luxCDABE* operon and MCS; $Amp^R$ | Karavolos *et al.* 2008 |
| pMK1*lux*-$P_{LEE1}$ | pMK1*lux* with TUV93-0 LEE1 promoter cloned into MCS; $Amp^R$ | This study |
| pMK1*lux*-$P_{aau}$ | pMK1*lux* with *Z0415* promoter cloned into MCS; $Amp^R$ | This study |
| pMK1*lux*-$P_{araB}$ | pMK1*lux* with TUV93-0 *araB* promoter cloned into MCS; $Amp^R$ | This study |
| pACYC184 | p15A ori multicopy plasmid; $Cm^R$, $Tet^R$ | Lab stock |
| pACYC184-*araC* | pACYC184 with TUV93-0 *araC* cloned into MCS; $Cm^R$, $Tet^R$ | This study |
| pSUPROM | Cloning vector for expression under the TatA promoter; $Kan^R$ | Jack *et al.* 2004 |
| pSUPROM-*araC* | pSUPROM with TUV93-0 *araC* cloned into MCS; $Kan^R$ | This study |
| pSUPROM-*araE* | pSUPROM with TUV93-0 *araE* cloned into MCS; $Kan^R$ | This study |
| pSUPROM-*araBADE* | pSUPROM with TUV93-0 *araBAD/araE* cloned into MCS; $Kan^R$ | This study |
| pKD46 | LRed recombinase expressing plasmid; temperature sensitive; $Amp^R$ | Datsenko and Wanner, 2000 |
| pKD3 | Template plasmid for LRed mutagenesis; $Cm^R$ | Datsenko and Wanner, 2000 |
| pKD4 | Template plasmid for LRed mutagenesis; $Kan^R$ | Datsenko and Wanner, 2000 |
| pCP20 | FLP recombinase expressing plasmid; temperature sensitive; $Amp^R$ | Datsenko and Wanner, 2000 |
| p*rpsM*:GFP | pACYC184, *rpsM*:GFP transcriptional fusion | Roe *et al.* 2004 |

**Supplementary Table 3** – Primers used in this study

| Primer name | Description | Sequence |
|---|---|---|
| *Z0415-9*_LRed_Fwd | *Z0415-9* KO forward primer | gcgcgctaatttgggcgaacacttcctgactaccctgcaatgaggctgaagtgtaggctggagctgcttc |
| *Z0415-9*_LRed_Rev | *Z0415-9* KO reverse primer | cgcctgatatgtcatcgcggcaaaacgcgtccattgaatatagccaatatcatatgaatatcctccttag |
| *Z0415-9*_Check_Fwd | *Z0415-9* KO forward check primer | tctctccagcgcgctaat |
| *Z0415-9*_Check_Rev | *Z0415-9* KO reverse check primer | atgtcatcgcggcaaaac |
| EHEC_*araE*_LRed_Fwd | EHEC *araE* KO forward primer | attgttcacgtatttttcactatgtcttactctctgctggcaggaaaaagtgtaggctggagctgcttc |
| EHEC_*araE*_LRed_Rev | EHEC *araE* KO reverse primer | ctctattaacgaaaaaagggccggatgtacagcacatccggcccgtgaaacatatgaatatcctccttag |
| EHEC_*araE*_Check_Fwd | EHEC *araE* KO check forward primer | aatatccatcacataacggcatg |
| EHEC_*araE*_Check_Rev | EHEC *araE* KO check reverse primer | attcccagctcattcctccc |
| EHEC_*araFGH*_LRed_Fwd | EHEC *araFGH* KO forward primer | ttttgccctgcacaaaacgacactaaagctggagagaaccgtgtaggctggagctgcttc |
| EHEC_*araFGH*_LRed_Rev | EHEC *araFGH* KO reverse primer | tgtggtgggaaaaaacgttaaattgttgtggaaaaaagcacatatgaatatcctccttag |
| EHEC_*araFGH*_Check_Fwd | EHEC *araFGH* KO check forward primer | tcccgctaaatttatgcacgt |
| EHEC_*araFGH*_Check_Rev | EHEC *araFGH* KO check reverse primer | ttgcaacgaagaacagccaa |
| EHEC_*araBAD*_LRed_Fwd | EHEC *araBAD* KO forward primer | gcaactctctactgtttctccatacccgtttttttggatggagtgaaacggtgtaggctggagctgcttc |
| EHEC_*araBAD*_LRed_Rev | EHEC *araBAD* KO reverse primer | aaaaaaccaggcttgattatagcctggtttcatttgattggctgtggttttatacagtcacatatgaatatcctccttag |
| EHEC_*araBAD*_Check_Fwd | EHEC *araBAD* KO check forward primer | cgtcacactttgctatgcca |
| EHEC_*araBAD*_Check_Rev | EHEC *araBAD* KO check reverse primer | aagataaaacctgcctgcgc |
| EHEC_*araC*_LRed_Fwd | EHEC *araC* KO forward primer | tgcaatatggacaattggtttcttctctgaatggcgggagtatgaaaagtgtgtaggctggagctgcttc |
| EHEC_*araC*_LRed_Rev | EHEC *araC* KO reverse primer | caaaccctatgctactccgtcaagccgtcaattgtctgattcgttaccaacatatgaatatcctccttag |
| EHEC_*araC*_Check_Fwd | EHEC *araC* KO check forward primer | tcttctctgaatggcgggag |

| | | |
|---|---|---|
| EHEC_*araC*_Check_Rev | EHEC *araC* KO check reverse primer | atggacgaagcagggattct |
| Crod_*araBAD*_LRed_Fwd | *C. rodentium araBAD* KO forward primer | cccactcactactgtttctccatacccgtatttctggatggagtgaaacggtgtaggctggagctgcttc |
| Crod_*araBAD*_LRed_Rev | *C. rodentium araBAD* KO reverse primer | tgtgttccggaataaaaatacgcgccactgtcgggacgcgtattttgcatcatatgaatatcctccttag |
| Crod_*araBAD*_Check_Fwd | *C. rodentium araBAD* KO check forward primer | acaacggcagaaatgtccac |
| Crod_*araBAD*_Check_Rev | *C. rodentium araBAD* KO check reverse primer | ctttcattcgctggagggc |
| pMK1*lux*-$P_{LEE1}$_EHEC_Fwd | Forward primer for cloning EHEC LEE1 promoter with EcoRI | cccgaattcctgtaactcgaattaagt |
| pMK1*lux*-$P_{LEE1}$_EHEC_Rev | Reverse primer for cloning EHEC LEE1 promoter with BamHI | cccggatccaatctccgcatgctttaata |
| pMK1*lux*-$P_{Z0415}$_Fwd | Forward primer for cloning *Z0415* promoter with EcoRI | cccgaattcattcaccagaaatggacg |
| pMK1*lux*-$P_{Z0415}$_Rev | Reverse primer for cloning *Z0415* promoter with BamHI | cccggatccatttcagcctcattgcag |
| pMK1*lux*-$P_{araB}$_Fwd | Forward primer for cloning *araB* promoter with EcoRI | cccgaattccgggaccaaagccatgac |
| pMK1*lux*-$P_{araB}$_Rev | Reverse primer for cloning *araB* promoter with XbaI | gcgctctagacgtttcactccatccaaa |
| pMK1*lux*_Check_Fwd | Forward primer to check pMK1*lux* cloning | ctataaaaataggcgtatcac |
| pMK1*lux*_Check_Rev | Reverse primer to check pMK1*lux* cloning | ctggccgttaataatgaatg |
| pACYC184-*araC*_Fwd | *araC* Gibson assembly forward primer | tgaagtcagccccatacgattgcaatcgccatcgtttca |
| pACYC184-*araC*_Rev | *araC* Gibson assembly reverse primer | caatccatgccaacccgttcttatgacaacttgacggct |
| pACYC184_Check_Fwd | Forward primer to check pACYC184 cloning | gacgctcaaatcagtggtgg |
| pACYC184_Check_Rev | Reverse primer to check pACYC184 cloning | gcattcacagttctccgcaa |
| pACYC184_Linear_Fwd | pACYC184 linearisation forward primer | gaacgggttggcatggattg |
| pACYC184_Linear_Rev | pACYC184 linearisation reverse primer | atcgtatggggctgacttca |
| pSUPROM-*araC*_Fwd | Forward primer for cloning *araC* with BamHI | ggccggatcctcttctctgaatggcgggag |
| pSUPROM-*araC*_Rev | Reverse primer for cloning *araC* with XbaI | ggcctctagaatggacgaagcagggattct |
| pSUPROM-*araE*_Fwd | Forward primer for cloning *araE* with BamHI | ggccggatcctgtcttactctctgctggca |

| | | |
|---|---|---|
| pSUPROM-*araE*_Rev | Reverse primer for cloning *araE* with XbaI | ggcctctagaaacgagacaaac gcctcaac |
| pSUPROM-*araBADE*_F1_Fwd | *araBAD* fragment Gibson assembly forward primer | tctaccacagaggaggatccatg gcgattgcaattggc |
| pSUPROM-*araBADE*_F1_Rev | *araBAD* fragment Gibson assembly forward primer | cagcagagagttactgcccgtaa tatgcc |
| pSUPROM-*araBADE*_F2_Fwd | *araE* fragment Gibson assembly forward primer | cgggcagtaactctctgctggcag gaaaaaatg |
| pSUPROM-*araBADE*_F2_Rev | *araE* fragment Gibson assembly forward primer | ctcgaggggtcgactctagatca gacgccgatatttctcaac |
| pSUPROM_Check_Fwd | Forward primer to check pSUPROM cloning | ctcttcgctattacgccagc |
| pSUPROM_Check_Rev | Reverse primer to check pSUPROM cloning | accctcatcagtgccaacat |
| pSUPROM_Linear_Fwd | pSUPROM linearisation forward primer | tctagactcgacccctcg |
| pSUPROM_Linear_Rev | pSUPROM linearisation reverse primer | ggatcctcctctgtggtag |
| *escT*_Fwd | *escT* RT-qPCR forward primer | tttgggctatagatgcggct |
| *escT*_Rev | *escT* RT-qPCR reverse primer | ggatgaatcgcttatagacggg |
| *escC*_Fwd | *escC* RT-qPCR forward primer | gctgaagtgagtgctcgttt |
| *escC*_Rev | *escC* RT-qPCR reverse primer | cctcaagcgggtcaataacg |
| *escV*_Fwd | *escV* RT-qPCR forward primer | ctaaaagttctccagtacgtgc |
| *escV*_Rev | *escV* RT-qPCR reverse primer | tcgccagagaaatcatcattca |
| *espA*_Fwd | *espA* RT-qPCR forward primer | ttcctgtaaatccgatgcgc |
| *espA*_Rev | *espA* RT-qPCR reverse primer | tggttgacgctttagatgcc |
| *tir*_Fwd | *tir* RT-qPCR forward primer | ttcctgtaaatccgatgcgc |
| *tir*_Rev | *tir* RT-qPCR reverse primer | atcgagcggaccatgatcat |
| *Z0415*_Fwd | *Z0415* RT-qPCR forward primer | tggtgtcttcgctgttattagg |
| *Z0415*_Rev | *Z0415* RT-qPCR reverse primer | cacggcataccatcgacttta |
| *Z0417*_Fwd | *Z0417* RT-qPCR forward primer | tggaagtttccgaccgtattt |
| *Z0417*_Rev | *Z0417* RT-qPCR reverse primer | tcatcaggaaaccgagttgtt |
| *Z0418*_Fwd | *Z0418* RT-qPCR forward primer | gccttactggttaatcgcctac |
| *Z0418*_Rev | *Z0418* RT-qPCR reverse primer | gtacagccacacacctttactc |
| Housekeeping_GroEL_Fwd | GroEL RT-qPCR forward primer | accgctgcagttgaagaa |
| Housekeeping_GroEL_Rev | GroEL RT-qPCR reverse primer | ctacggtttcgtcggagttag |
| Housekeeping_GapA_Fwd | GapA RT-qPCR forward primer | cggtaccgttgaagtgaaaga |
| Housekeeping_GapA_Rev | GapA RT-qPCR reverse primer | acttcgtcccatttcaggttag |
